# Predicting fine-scale distributions and emergent spatiotemporal patterns from temporally dynamic step selection simulations

**DOI:** 10.1101/2024.03.19.585696

**Authors:** Scott W Forrest, Dan Pagendam, Michael Bode, Christopher Drovandi, Jonathan R Potts, Justin Perry, Eric Vanderduys, Andrew J Hoskins

**Affiliations:** School of Mathematical Sciences, Queensland University of Technology, Australia; Centre for Data Science, Queensland University of Technology, Australia; CSIRO Environment, Dutton Park, Australia; CSIRO Data61, Dutton Park, Australia; School of Mathematics and Statistics, University of Sheffield, Sheffield, UK; Northern Australian Indigenous Land and Sea Management Alliance, Darwin, Australia; CSIRO Environment, Townsville, Australia

**Keywords:** animal movement, circadian, fine-scale dynamics, harmonics, landscape-scale distributions, simulated trajectories, step selection functions, temporal dynamics

## Abstract

Understanding and predicting animal movement is fundamental to ecology and conservation management. Models that estimate and then predict animal movement and habitat selection parameters underpin diverse conservation applications, from mitigating invasive species spread to enhancing landscape connectivity. However, many predictive models overlook fine-scale temporal dynamics within their predictions, despite animals often displaying fine-scale behavioural variability that might significantly alter their movement, habitat selection and distribution over time. Incorporating fine-scale temporal dynamics, such as circadian rhythms, within predictive models might reduce the averaging out of such behaviours, thereby enhancing our ability to make predictions in both the short and long term. We tested whether the inclusion of fine-scale temporal dynamics improved both fine-scale (hourly) and long-term (seasonal) spatial predictions for a significant invasive species of Northern Australia, the water buffalo (*Bubalus bubalis*). Water buffalo require intensive management actions over vast, remote areas and display distinct circadian rhythms linked to habitat use. To inform management operations we generated hourly and dry season prediction maps by simulating trajectories from static and temporally dynamic step selection functions (SSFs) that were fitted to the GPS data of 13 water buffalo. We found that simulations generated from temporally dynamic models replicated the buffalo’s crepuscular movement patterns and dynamic habitat selection, resulting in more informative and accurate hourly predictions. Additionally, when the simulations were aggregated into long-term predictions, the dynamic models were more accurate and better able to highlight areas of concentrated habitat use that might indicate high-risk areas for environmental damage. Our findings emphasise the importance of incorporating fine-scale temporal dynamics in predictive models for species with clear dynamic behavioural patterns. By integrating temporally dynamic processes into animal movement trajectories, we demonstrate an approach that can enhance conservation management strategies and deepen our understanding of ecological and behavioural patterns across multiple timescales.

## 1 Introduction

The movement of animals through space and time is critical to ecosystem functioning. Predicting the movement and distribution of animals across a landscape is difficult, however, as the successive fine-scale behavioural decisions that animals make are complex and rarely match the scale of our observations or the target of our predictions (Levin, 1992). These complex decision-making processes relate to extrinsic factors such as resources, weather and other animals, and intrinsic factors such as age, sex, memory and energetic state (Turchin, 1998; Nathan et al., 2008; Morales et al., 2010; Fagan et al., 2013). These movement and habitat selection behaviours also change throughout time across various scales, which often relate to natural cycles. Animals typically have daily rhythms that mediate foraging or hunting behaviours, resting, thermoregulation and finding water (Ouled-Cheikh et al., 2020; Richter et al., 2020; Toro-Cardona et al., 2023), and often have seasonal rhythms determined by the climate, availability of resources, and breeding behaviour (Tulloch, 1970; Ager et al., 2003; Rafiq et al., 2023).

There are countless examples of diel cycles (periodic over 24 hours) in the animal kingdom, and these rhythms are often defined by significant changes in movement and habitat selection behaviour (Fryxell et al., 2008; Webb et al., 2010; Buderman et al., 2018; Thaker et al., 2019; Meese & Lowe, 2020; Richter et al., 2020; Nisi et al., 2022). The mention of a diurnal, crepuscular or nocturnal species evokes substantial diel behavioural changes, and in some cases animals show opposing trends between different periods throughout the daily period, such as an animal that needs to forage in open areas but seeks shelter when it is vulnerable to predation (Leblond et al., 2010; Kohl et al., 2018; Richter et al., 2020; Palmer et al., 2022). In many fishes, diel patterns of activity are common, and a range of diurnal, nocturnal and crepuscular foraging strategies define niche segmentation, as there are specific adaptations that are beneficial to each strategy (Fox & Bellwood, 2011; Currey et al., 2015).

Despite our understanding that animal behaviour is temporally variable on fine scales, these dynamics are often overlooked when developing predictive models. We pose that including them can allow for two-fold benefits. Firstly, including fine-scale temporal dynamics allows for temporally fine-scale predictions. There are many examples where temporally fine-scale predictions would be valuable, such as for predicting human-wildlife conflict and poaching (Carter et al., 2012; Buderman et al., 2018; Forrest et al., 2024), identifying high-risk locations and times for zoonotic disease transfer (Parsons et al., 2014), mitigating the impacts of fisheries on wildlife (Ferńandez & Anderson, 2000; Ouled-Cheikh et al., 2020), and for optimising intensive conservation management actions such as trapping, shooting and mustering. Secondly, including fine-scale temporal dynamics has the potential to provide more accurate long-term predictions, as the dynamic processes of movement and habitat selection are explicitly represented rather than being averaged out. An animal that selects between open and closed vegetation at different times of the day (Leblond et al., 2010; Kohl et al., 2018) will have qualitatively different patterns of space use than a species that selects for medium canopy cover at all times, but ignoring the dynamic behaviour will result in the average effect (selection for medium canopy), leading to different and possibly less accurate predictions. To the best of our knowledge, assessing whether fine-scale temporally dynamic processes lead to more accurate long-term predictions has not been formally tested in the literature.

As the distribution of animals from one hour to the next is correlated due to the animal’s movement dynamics, temporally fine-scale predictions can be difficult to generate analytically. However, a flexible and robust approach to generating temporally fine-scale predictions is to simulate stochastic animal movement trajectories (Morales et al., 2010; Osipova et al., 2019; Hooker et al., 2021; Whittington et al., 2022; Aiello et al., 2023; Hofmann et al., 2023; Potts & Börger, 2023; Sells et al., 2023). Simulation-based models can include temporal dynamics on any scale, and they explicitly incorporate the animal’s movement dynamics. Step selection functions (SSFs) are particularly advantageous as they can be used to simulate trajectories (Signer et al., 2017; Potts & Börger, 2023; Signer et al., 2023), are straightforward to parameterise, and can incorporate temporal dynamics (Ager et al., 2003; Forester et al., 2009; Tsalyuk et al., 2019; Richter et al., 2020; Klappstein et al., 2024). An SSF combines a movement and a external selection kernel, can take a range of forms (Munden et al., 2021; Klappstein et al., 2022; Beumer et al., 2023; Eisaguirre et al., 2024; Pohle et al., 2024), and can accommodate a wide range of covariates including habitat, linear features, distance-to-feature variables, proximity to other animals (Potts et al., 2022; Ellison et al., 2024) and representations of previous space use (Schlägel & Lewis, 2014; Oliveira-Santos et al., 2016; Rheault et al., 2021).

The water buffalo (Bubalus bubalis) population in Northern Australia’s tropical savannas requires accurate spatiotemporal predictions on both fine scales and longer periods for managers to make informed conservation decisions (Werner, 2005; Mihailou & Massaro, 2021). Globally, tropical savanna ecosystems are undergoing significant change and degradation (Williams et al., 2022), and in Northern Australia, feral animals such as pigs (*Sus scrofa*), cattle (predominately *Bos indicus*) and buffalo are a major destructive force, causing significant environmental damage and presenting a major disease threat to Australia’s agricultural systems (e.g. brucellosis, tuberculosis and African swine fever) (Skeat et al., 1996; Werner, 2005; Petty et al., 2007; Mihailou & Massaro, 2021). Management of these feral vertebrates is usually carried out using costly and intensive mustering or culling operations. With accurate predictions of distribution at temporally fine-scales, conservation managers can more effectively target control measures, with the potential to simulate alternative scenarios (Warwick-Evans et al., 2018). With accurate long-term predictions, areas that are most at-risk can be identified by high predicted buffalo use, allowing for persistent management operations such as exclusion fencing to be considered (Ens et al., 2016; Sloane et al., 2024).

To assess the benefits of including fine-scale temporal dynamics in predictive models, our objectives in this paper are to (1) investigate whether incorporating fine-scale temporal dynamics into SSFs can be used to generate accurate hourly predictions via simulated trajectories, and (2) whether predictions over the longer time-scales reveal emergent features that are captured only with dynamic models, and provide higher prediction accuracy than models that only incorporate static parameters. To assess these objectives, we simulated models that include circadian temporal dynamics with varying flexibility, which were fitted to and validated against hourly GPS tracking data of buffalo. We hypothesised that the predictions on the hourly scale would be more infor-mative, as the behaviour of buffalo is highly variable throughout the day in Northern Australia’s tropical savannas, and that the long-term predictions will more closely represent the distribution of the observed data, as there are landscape features that buffalo use that will be ignored when using a static model that averages over movement and habitat selection behaviour.

## 2 Methods

### 2.1 Study area and data collection

Data were collected from the Djelk Indigenous Protected in Western Arnhem Land, Northern Territory, Australia. The area is a culturally significant landscape comprised of tropical savanna with areas of open woodland, rainforest, a varied river and wetland system, and open floodplains. To understand their fine-scale and long-term movement and habitat selection behaviours, water buffalo (*Bubalus bubalis*) were GPS-tracked in collaboration with Djelk rangers between July 2018 and November 2019. Due to their size and potential risk to handlers, wild buffalo were immobilised via a helicopter using tranquiliser darts following a modified version of the methods described in McMahon and Bradshaw (2008). All anaesthesia procedures on buffalo were undertaken by a qualified wildlife veterinarian. Animals were equipped with a device that was a combination of a commercial VHF tracking collar (Telonics MOD-500) and a custom developed LoRa (long-range) radio enabled GPS tracking pod. In total the VHF collar and LoRa-GPS pod weighed 1.2 kg. The LoRa-GPS pods were scheduled to collect a GPS location once an hour with an accuracy of 10 m, and attempted to transmit GPS locations over a custom-installed LoRa network. GPS locations were also logged onboard the device, which is the data that was used in this study. In total 17 female buffalo were GPS-tracked, all devices were retrieved, and of those we used 13 individuals that had high-quality data with temporally consistent fixes for at least 3 months. The hourly GPS locations had a fix-success rate of 88% on average (range = 59% - 95%).

### 2.2 Landscape covariates

Buffalo’s movement decisions are driven by factors such as vegetation composition and density for resource acquisition and shade, access to water, and the terrain (Campbell et al., 2020). In monsoonal ecosystems of Northern Australia, vegetation and the distribution of water changes dramatically throughout the year. To represent the seasonal changes in vegetation, we used monthly Normalised Difference Vegetation Index (NDVI), which was derived from Sentinel-2 remote sensing data and measures photosynthetic activity and approximates the density and health of vegetation (Reed et al., 1994; Myneni et al., 1995). NDVI is an informative covariate in this landscape (Campbell et al., 2020) as it distinguishes between the broad vegetation classes, identifies wet and flooded areas, and can quantify buffalo’s forage resources as they are typically under open canopy. Monthly NDVI layers were generated from Sentinel-2 spectral imagery at 10 m x 10 m resolution using Google Earth Engine by taking the clearest pixels from a range of images for that month to alleviate the effects of obstruction from clouds. We selected the highest quality-band-score, which is based on cloud and shadow probability for each pixel, resulting in a single obstruction-free image of the NDVI values for each month of each year. We also used temporally-static layers for canopy cover and herbaceous vegetation, which are derivatives of Landsat-7 imagery and were sourced from Geoscience Australia at 25 m x 25 m resolution (Source: Geoscience Australia; Landsat-7 image courtesy of the U.S. Geological Survey). We represented the terrain by including a slope covariate, which was summarised from a 25 m x 25 m digital elevation model using the terra R package (Hijmans, 2024; R Core Team, 2024), and was calculated using the methodology of Horn (1981). The canopy cover layer was a proportion from 0 (completely open) to 1 (completely closed), and the herbaceous vegetation layer was binary, with 1 representing grasses and forbs, and 0 representing other (which is predominately woody growth). All spatial variables were discretised into grids (i.e. rasters) and resampled to be 25 m x 25 m resolution.

### 2.3 SSF model and temporally-varying parameters

Step selection functions (SSFs) are statistical models commonly applied to telemetry data to infer movement and habitat selection behaviour (Fortin et al., 2005; Potts et al., 2014; Thurfjell et al., 2014; Avgar et al., 2016; Northrup et al., 2022; Potts & Börger, 2023; Michelot et al., 2024). To denote the SSF, *p* is the likelihood of observing an animal at location *s_t_* given its last two observed locations *s_t_*_−1_, *s_t_*_−2_ with locations indexed by discrete time *t* and within a spatial domain *S*. Let **X** be a multivariate spatial field indexed by a location *s_t_*, and where **X**(*s_t_*) = (*X*_1_(*s_t_*)*, …, X_n_*(*s_t_*)) is a vector with a length equal to the number of spatial covariates:

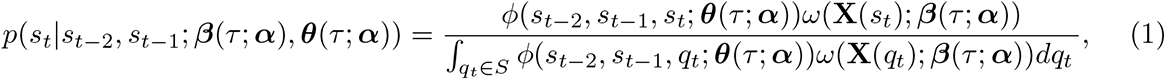

Here, *ϕ* denotes the movement kernel, with parameters relating to step lengths and turning angles contained in ***θ***(*τ*; ***α***), which may be some function of time which we denote here as *τ*, as this time component may be cyclic and therefore not be equal to the absolute time, *t*. The parameter vector ***α*** defines the functional relationship between ***θ*** and *τ*, as well as between ***β*** and *τ* in the *ω* component. The *ω* component is the external selection kernel, and the term in the denominator ensures that the probability density, *p*(·), integrates to 1, which in practice is typically approximated through numerical integration, where we sample a set of proposed next steps given what is ‘available’ to the animal as determined by the movement kernel (Avgar et al., 2016; Michelot et al., 2024).

The external selection term is typically modelled analogously to a resource-selection function (RSF), that assumes an exponential (log-linear) form as

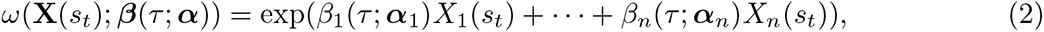

where ***β***(*τ*; ***α***) = (*β*_1_(*τ*; ***α***_1_), . . ., *β*_*n*_(*τ*; ***α***_*n*_)),

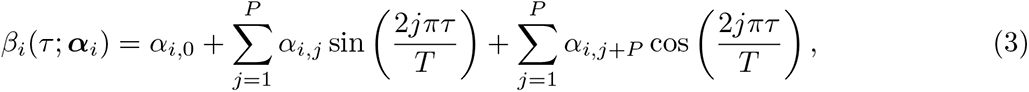

and ***α****_i_* = (*α_i,_*_0_*, …, α_i,_*_2_*_P_*). Here, we have used harmonic terms to define the functional relationship between *τ* and ***β*** (Ager et al., 2003; Forester et al., 2009; Tsalyuk et al., 2019; Richter et al., 2020). These are pairs of sine and cosine terms that have cyclic periods of varying frequency, such as sin(2*πτ/T*) and cos(2*πτ/T*), where *τ* ∈ *T*. Harmonics are a natural choice as they are cyclic, which often aligns with temporal changes that animals respond to, such as daily or seasonal cycles (Boyce et al., 2010), although other additive terms such as splines are another obvious choice (Hanks et al., 2015; Klappstein et al., 2024). For a yearly cycle for instance, *τ* would represent the day of the year, and *T* would equal 365, although *τ* does not need to be integer-valued and can be arbitrarily fine.

### 2.4 SSF data preparation and model fitting

Buffalo have temporally dynamic patterns in their movement and external selection behaviours across two predominant time-scales: daily and seasonal. Daily rhythms are predominately driven by daylight and temperature, and seasonal rhythms are predominately driven by the distribution of water and the availability of forage (Tulloch, 1970; Campbell et al., 2020). As we wanted to generate temporally fine-scale distributions we included harmonic terms that have a daily periodicity. As buffalo management actions such as mustering predominately take place during the dry season because the landscape is more accessible, we separated data into wet and dry seasons and focused on dry season models with fine-scale temporal dynamics. An extension of this work might be to incorporate multiple interacting time-scales, such as daily rhythms that also change across the seasons.

To incorporate the temporal dynamics on a daily time-scale, each movement parameter and environmental covariate was included in the model as a linear predictor as well as interacting with pairs of harmonic terms relating to the hour of the day (Forester et al., 2009; Street et al., 2016; Tsalyuk et al., 2019; Warton, 2022). The resulting set of harmonics that we selected from to fit models was sin(2*kπτ/*24) and cos(2*kπτ/*24) for *k* = 1, 2, 3, where *τ* is indexed by the hour of the day, resulting in four models with 0, 1, 2 and 3 pairs of sine and cosine terms. Each movement parameter and spatial covariate therefore has up to seven estimated parameters, which can be combined into a single function that varies across the day. To allow for external selection to operate over a region of the environmental space (rather than monotonically increasing or decreasing), we also included quadratic parameters for NDVI (i.e. NDVI + NDVI^2^) and canopy cover, which were also interacted with the harmonic terms. The final set of models that were fitted to the dry season buffalo data were models with 0 pairs of harmonics (denoted 0p, or the static or ‘daily average’ model), and models with 1, 2 and 3 pairs of harmonics (dynamic models), which we denoted as 1p, 2p and 3p, respectively.

As we were fitting a model across all individuals simultaneously, we generated ‘available’ steps using the amt package (Signer et al., 2019) by fitting a gamma distribution to the observed step lengths and a von Mises distribution to the observed turning angles of all individuals, which makes ‘updating’ the movement parameters after fitting the SSF straightforward (Fieberg et al., 2021). We sampled independently from these distributions to generate new steps, and matched each observed step with 10 randomly-generated available steps.

Due to the large number of covariates when including several pairs of harmonics, particularly when also including quadratic terms, we had difficulty fitting population-level models with the ‘glmmTMB’ (Brooks et al., 2017) and ‘INLA’ (Rue et al., 2009) model fitting approaches (Muff et al., 2020). We therefore fitted models to all individuals using the ‘TwoStep’ approach from the TwoStepCLogit package (Craiu et al., 2011, 2016) in R (R Core Team, 2024), which can be considered as a more computationally efficient alternative to fitting models with individual-level random effects, providing certain conditions are met, namely that all individuals visit every category level when using categorical covariates (Muff et al., 2020). The first step of the TwoStep approach is to fit individual-level conditional logistic regression models, which are then combined in the second step using the expectation-maximisation (EM) algorithm in conjunction with conditional restricted maximum likelihood to estimate the population-level parameters (Craiu et al., 2011). For computational stability, prior to model fitting we centred and scaled all variables by subtracting mean values and dividing by the standard deviation, using the data from all used and random steps. The centering and scaling was done immediately prior to model fitting, and therefore after creating the quadratic terms and multiplying the covariates with the harmonics. Following model fitting, the coefficients and covariates were returned to their natural scale for interpretation and prediction, and the movement parameters (shape and scale of the gamma distribution and concentration parameter of the von Mises distribution) were updated across *τ* following Avgar et al. (2016) and Fieberg et al. (2021).

### 2.5 Generating trajectories from the fitted model

To generate trajectories, we first pick a starting location uniform at random from the study domain. Then *n* steps are drawn from the ‘updated’ movement kernel (Fieberg et al., 2021) by sampling independently from the updated gamma and von Mises distributions of step-lengths and turning angles for time *τ*, respectively. For each proposed step, exp (***β***(*τ*; ***α***)**X**(*s_t_*)) (Equation 2) is evaluated, resulting in a value for each proposed step that is proportional to the probability of being selected, and a single step is probabilistically selected based on these values. This new step is added to the trajectory and becomes the starting point for the next step, and the process is repeated until the specified number of steps are reached. For our simulated trajectories we proposed 50 steps from the movement kernel at each time point.

### 2.6 Landscape-scale predictions

To estimate the expected distribution of buffalo predicted by the models for each hour of the day and for the late dry season, we simulated 100,000 trajectories with 3,000 steps each from the coefficients estimated by the four models (0p, 1p, 2p and 3p). We assessed the convergence of the simulations by randomly sampling 10 subsets of increasing numbers of trajectories (without replacement), and calculating the standard deviation of normalised (from 0 to 1) prediction values in 1000 randomly distributed but consistent cells (Figure A3). Each simulated trajectory took approximately 1 minute to run, which can easily be parallelised as each simulation is independent, and which we ran on a high performance computing cluster. As the landscape in which we collected our data was large (c. 60 km x 60 km) and mostly without observed buffalo GPS locations, we selected a 20 km x 20 km subset that contained a high density of buffalo locations for the late dry season 2018 (n = 13,940) from 8 individuals (mean ± SD = 1742 ± 596 GPS locations per buffalo). The covariates included in the simulations were the four spatial layers used to fit the models (average NDVI for the late dry season 2018, canopy cover, herbaceous vegetation and slope) at 25m x 25m resolution. To create the landscape-scale maps of expected buffalo distribution, we overlaid a template raster layer with 50 m x 50 m cells and summed the number of simulated locations that fell within each cell. For the long-term distribution maps all locations were considered for each model, and for the hourly distribution maps we separated the simulated locations into the respective hours and summed the number of simulated locations that fell within each template raster cell for each hour. We considered the number of simulated locations within each cell to approximate the expected spatial distribution of buffalo that was predicted by each model.

### 2.7 Comparing models and validating predictions

We assessed whether the simulations could recreate the daily behavioural dynamics of the water buffalo by summarising movement and habitat selection behaviour for each hour of the day. To achieve this we binned the steps of both the observed and simulated trajectories into the hours of the day, and took the mean, median and standard deviation of step lengths and the four habitat covariates - NDVI, canopy cover, herbaceous vegetation and slope, and visually compared across the hours of the day.

To assess the accuracy of the predicted distribution maps we used the continuous Boyce index (cBI) (Boyce et al., 2002; Hirzel et al., 2006). The cBI uses a sweeping window that separates habitat suitability (in our case number of simulated locations per cell) into bins, and assesses how many observed locations fell into these cells relative to their prevalence in the landscape, denoted as the F-ratio. For simulated locations that were completely random within a particular range of habitat suitability the F-ratio would be 1, indicating that cells are used in proportion to their prevalence in the landscape. For an accurate model, we would expect fewer true locations in areas that are predicted to be unsuitable habitat (resulting in an F-ratio less than 1), and more true locations in areas that are predicted to be more favourable habitat (resulting in an F-ratio greater than 1). This provides an F-ratio curve which is used to assess the performance of the model. We assessed whether there was an increasing trend of the F-ratio curve by using the Spearman rank correlation coefficient (hereafter Spearman correlation or *ρ*), which assesses monotonicity and ranges from −1 to 1, with perfect (monotonically increasing) predictions resulting in a value of 1 (Hirzel et al., 2006). A high value of *ρ* would indicate the the true GPS locations were observed in a similar pattern of use to what was predicted by the model. For the long-term distribution maps all 13,940 observed buffalo GPS locations that were within the landscape were used for validation, resulting in a single F-ratio curve and *ρ* per model. For the hourly distribution maps we used the observed buffalo locations for that hour (mean ± SD = 580 ± 68 locations per hour) to validate the predictions, resulting in a F-ratio curve and *ρ* for each hour of the day for each model.

## 3 Results

For the dynamic models that included harmonic terms there was clear temporal variability throughout the day for the buffalo’s movement and external selection parameters, in both the dry and wet seasons (Figure 2 and 3, and **??**). The external selection parameters indicated an avoidance of herbaceous vegetation (Figure 2) and selection of higher values of NDVI and canopy cover during the middle of the day, which correlates with buffalo thermoregulation as they seek shelter from high temperatures and sun (Figure 3). There was also positive selection for herbaceous vegetation in the early morning and evening, which are likely to represent foraging periods (Figure 2). When the harmonic terms are not considered, the dynamics in relation to herbaceous vegetation average to a coefficient almost equal to 0, obscuring any relationship to herbaceous vegetation.

**Figure 1:**
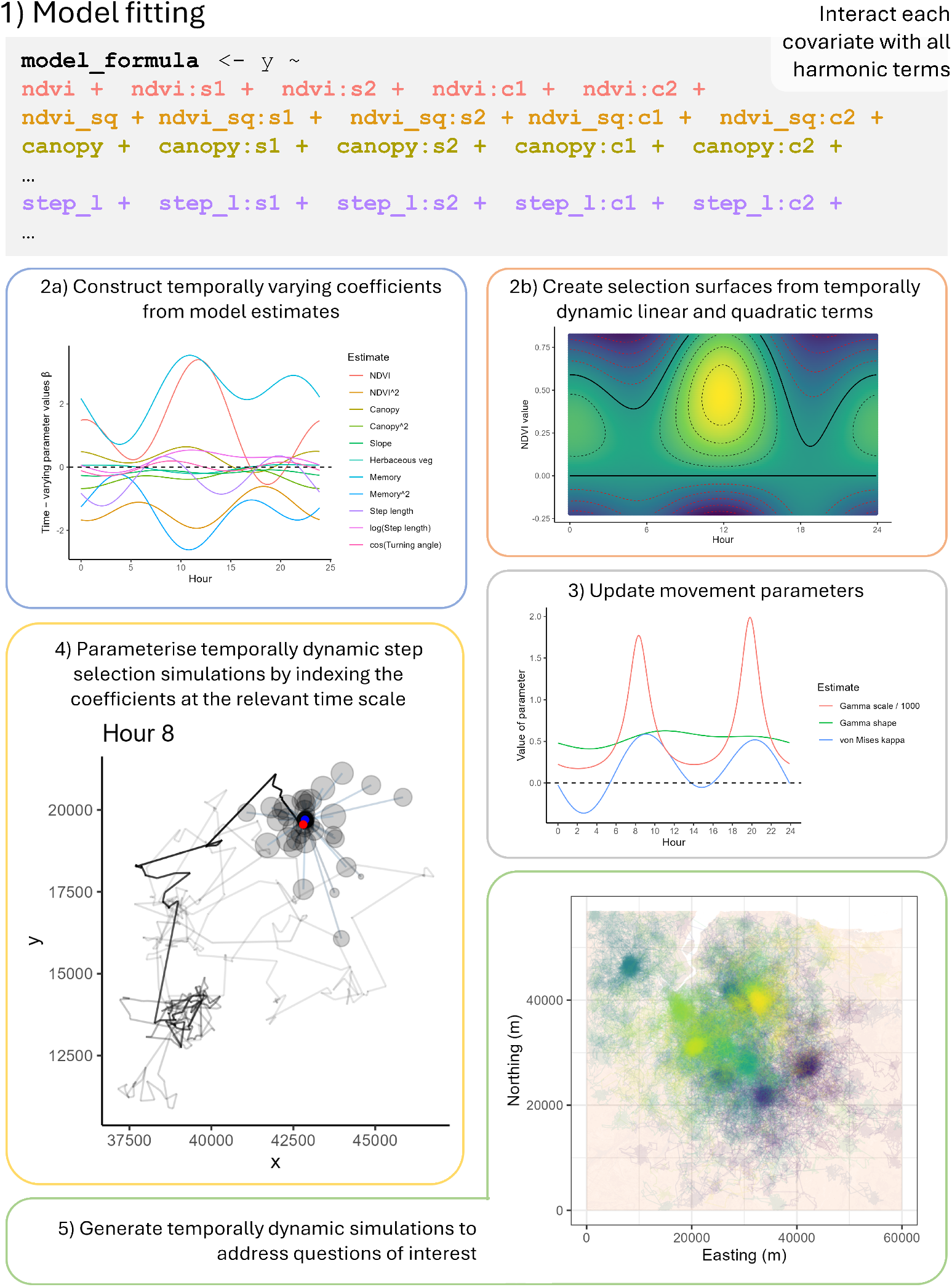
Concept plot illustrating the procedure of model fitting with harmonic terms, reconstructing the temporally dynamic coefficients, and generating temporally dynamic trajectories. The model is fitted with covariates interacting with the harmonic terms, where *s*1, *s*2, *c*1 and *c*2 in the model formula represent sin(2*πτ/*24), sin(4*πτ/*24), cos(2*πτ/*24) and cos(4*πτ/*24), respectively, and *τ* is the hour of the day where *τ* ∊ *T*, with *T* denoting 24 hours. All covariates can be interacted with the harmonics, including quadratic terms, movement parameters, memory and social covariates. Here we show simulations with a memory component **??**. The coefficients are then reconstructed into time-varying coefficients (2a), and when linear and quadratic terms are interacted with harmonic terms, ‘selection surfaces’ can be created (2b). The movement parameters are ‘updated’ following the typical procedure (Avgar et al., 2016; Fieberg et al., 2021), although this is performed across *T* (3). In our case there were negative values for the von Mises concentration parameter, suggesting that the mean of the distribution changed from 0 to *π*, indicating a higher likelihood of taking return steps in the early morning. Using the temporally dynamic external selection coefficients and updated movement parameters, we can simulate by indexing the coefficients and parameters at time *τ* (4). In (4), we have shown the proposed steps from the current (blue) point, with the size of the circle representing the probability of choosing that step based on the habitat covariates and previous locations. The red circle was the step that was selected, and forms the starting point for the next step.

**Figure 2:**
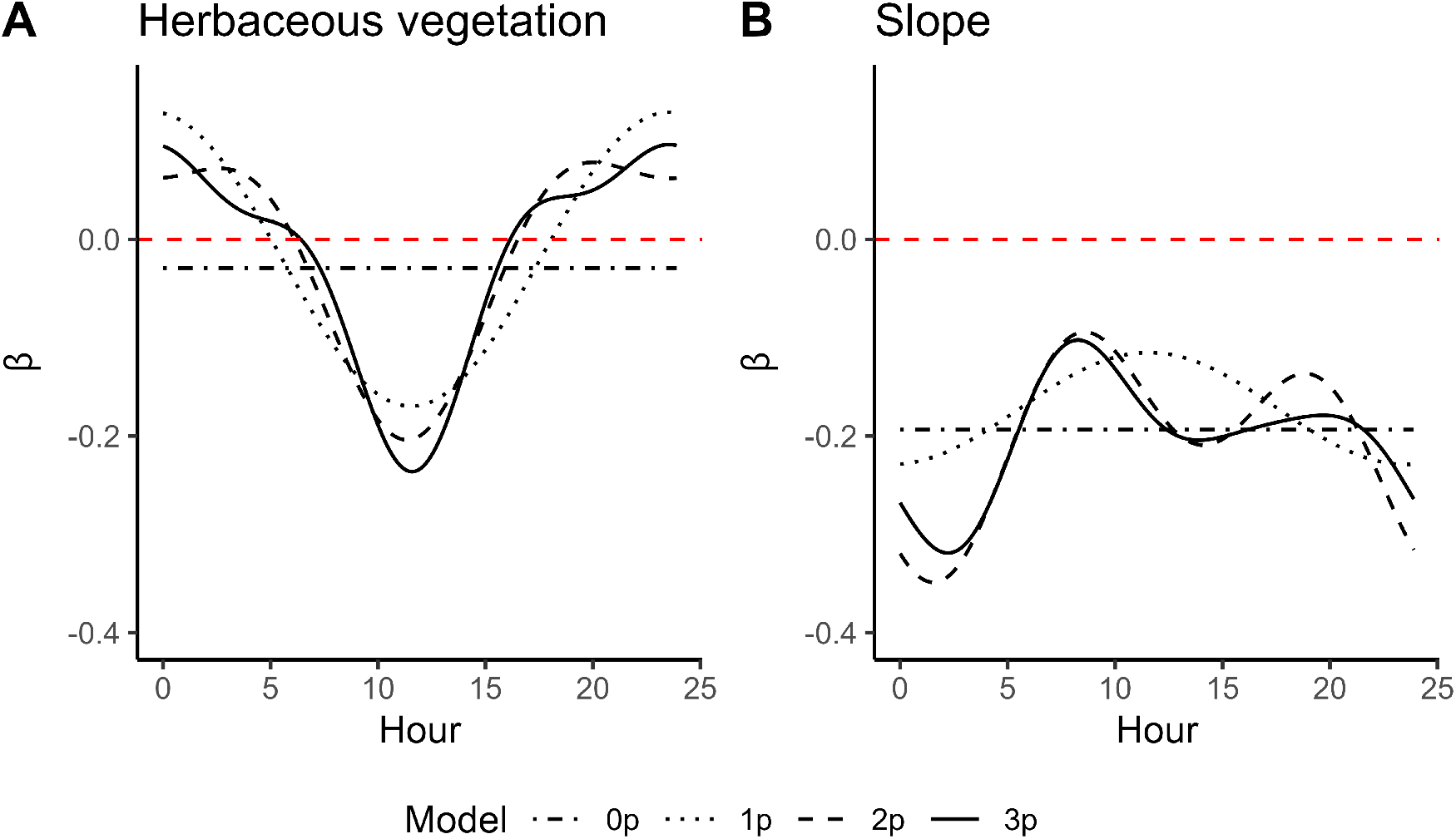
Estimated time-varying coefficients over the day for the external selection parameters of the population-level SSF model for 13 female buffalo in Arnhem Land, NT. Panel A shows the estimated coefficient for herbaceous vegetation for the four models. The static (0p) model averages over the effect throughout the day, suggesting incorrectly that there is little relationship to herbaceous vegetation. The dynamic models, however, reveal that there is attraction in the evening and avoidance during the day, suggesting buffalo seek shelter from the sun in woody vegetation during the middle of the day. As there appears to be only a single period in this trend, all three dynamic models (1p, 2p and 3p) capture it. The trend for slope appears to have multiple modes, which is captured by the models with two and three pairs of harmonics.

**Figure 3:**
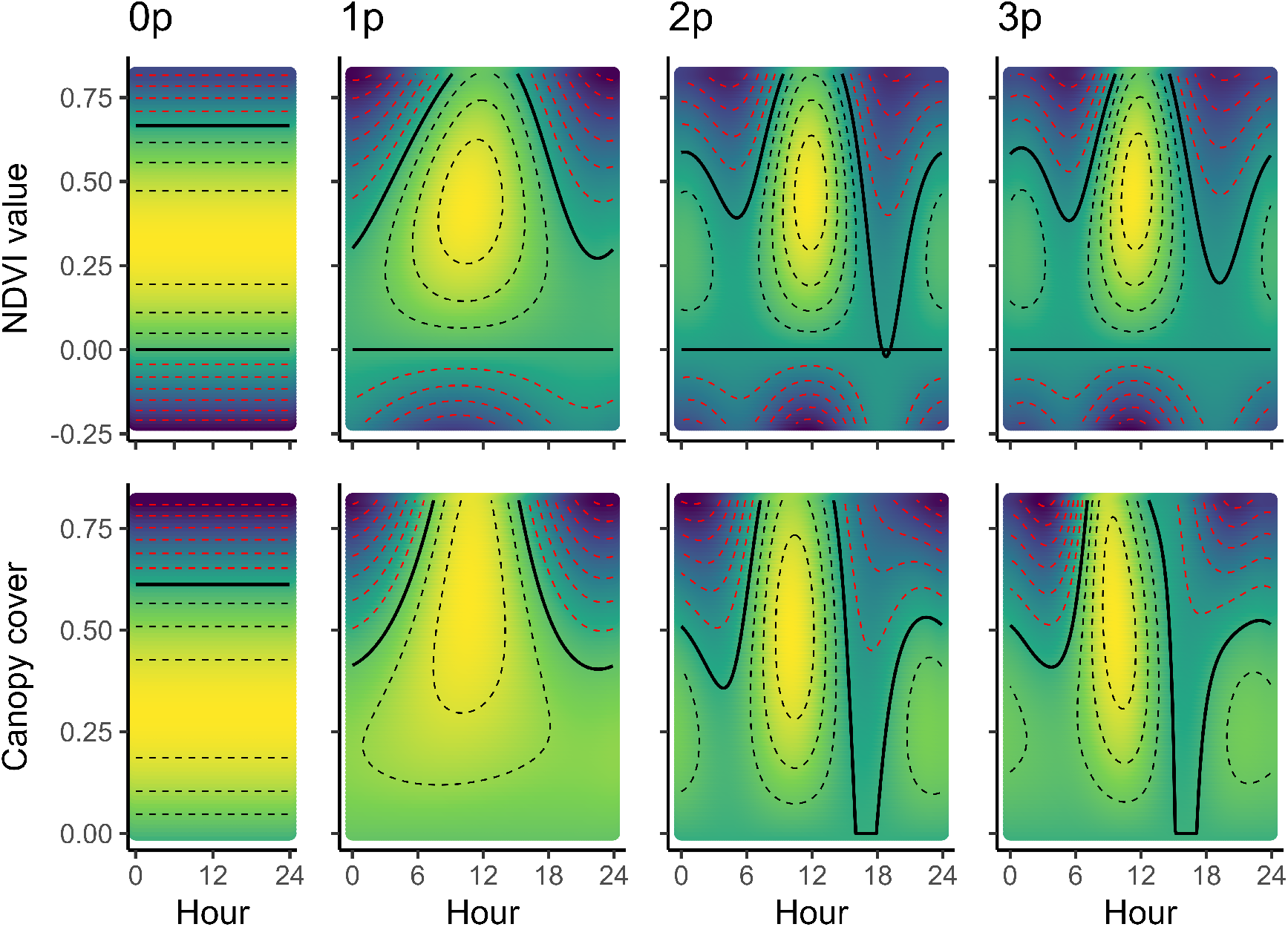
As there were both linear and quadratic terms that interacted with the harmonic terms for Normalised Difference Vegetation Index (NDVI) and canopy cover, it is possible to construct ‘selection surfaces’ that represent the strength of association through time on the natural scale of the covariate. The colours approaching yellow with the black dashed lines indicate positive selection with an estimated *β* coefficient above 0, the solid black line shows the zero contour, and colours approaching purple with red dashed contours showing avoidance of those values of each covariate at those times. Due to the difference in scales of the covariates, the selections colours are scaled for each plot. Each column represents a model with numbers of harmonics that increase to the right (0p, 1p, 2p and 3p denote 0, 1, 2 and 3 pairs of harmonics, respectively). Each row represents a different covariate within that model. For NDVI (top row) and canopy cover (second row), the models with one, two and three pairs of harmonics (1p, 2p and 3p) are similar, suggesting that increasing the number of harmonics is unlikely to dramatically change the model fit and therefore simulation outputs. The NDVI selection surface suggests that buffalo have an attraction for intermediate values of NDVI in the middle of the day, and selection against high values in the dawn and dusk periods, possibly for foraging and to ease transit in high movement periods. The selection surface for canopy cover indicates that buffalo prefer denser canopy in the middle of the day (correlating with NDVI), suggesting that buffalo are seeking refuge for high temperatures and sun.

The buffalo’s observed movement behaviour followed a crepuscular pattern, with high movement around dawn and dusk, and the simulated behaviour from the dynamic models with 2 or 3 pairs of harmonics (2p and 3p) replicated the pattern that was observed in the data (Figure 4), which was suggested by the estimated coefficients (Figure A1). The simulations from the dynamic 1p, 2p and 3p models largely capture the habitat selection during the middle of the day (Figure 4), but the selection of NDVI and canopy cover outside of the middle of the day were not well captured by any of the models.

**Figure 4:**
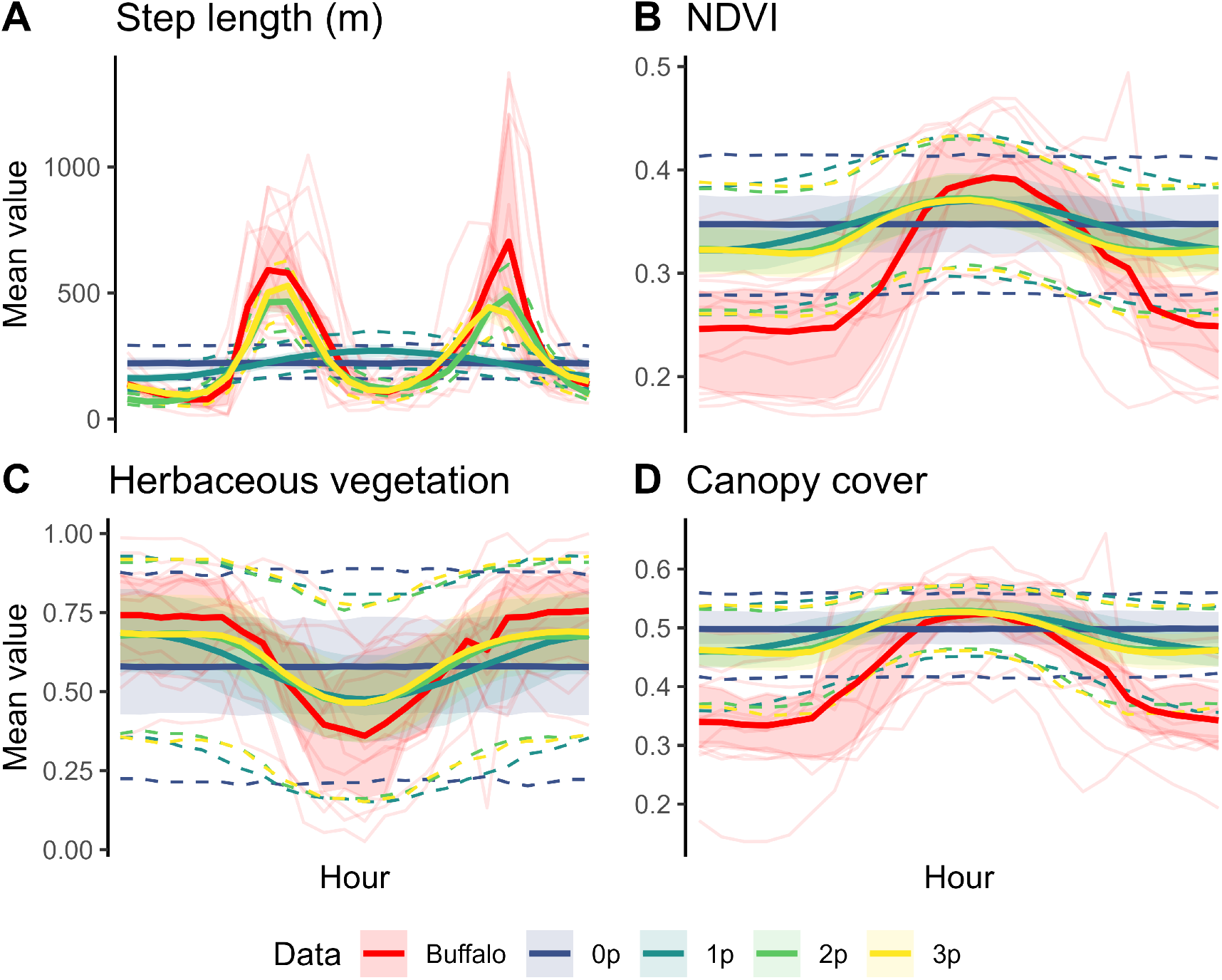
To assess the habitat selection of the simulated trajectories through time, we binned 100 randomly-selected trajectories into the hours of the day, extracted the values of the movement parameters and covariates for each hour, and took the mean, median and standard deviation. Here we show the mean values for the observed data and the simulations from each of the four models through time for (A) step length, (B) Normalised Difference Vegetation Index (NDVI), (C) herbaceous vegetation and (D) canopy cover. The shaded ribbons enclose the 25% and 75% quantiles, and the dashed lines are the 2.5% and 97.5% quantiles. The solid line is the mean for that hour across all trajectories for the buffalo or for each of the models. The observed data is shown in red, with each light red line representing one buffalo (*n* = 13). The dashed lines for the 2.5% and 97.5% quantiles are not displayed for the observed data, as all buffalo individuals are shown. For step length in Panel A, the simulated trajectories of the step selection models with two or three pairs of harmonics (2p and 3p) captured the bimodal, crepuscular movement pattern in the observed buffalo data. For the external covariates NDVI, herbaceous vegetation and canopy cover in Panels B-D, the simulated trajectories of the temporally dynamic models with one, two or three pairs of harmonics (1p, 2p, and 3p) largely capture the peak of the unimodal pattern of habitat selection in the dry season, which is driven by sun and high temperatures during the middle of the day, leading to seeking shelter in denser, more closed vegetation. Although the general shape is represented, none of the models capture the selection of low values of NDVI and canopy cover during the early morning and night, which is borne out in the validation of the landscape-scale predictions, which receive lower scores during these periods. The pattern of step lengths and habitat selection is consistent across the buffalo and within each model’s simulations, although the variance is wide, likely due to the area of the landscape that is available to the buffalo or the simulated trajectories.

When considering the long-term landscape-scale predictions, the static (0p) model’s predictions are more homogeneous, indicating an averaging-out of high habitat selection, such as during the middle of the day (Figure 5). The predictions of the most accurate dynamic (2p) model on the other hand, highlight areas of concentrated use in the landscape, which aligned better with the observed buffalo locations (Figure 5). The continuous Boyce index (cBI) indicated that the 2p model was most accurate (*ρ* = 0.92), followed by the 3p (*ρ* = 0.82), 0p (*ρ* = 0.70) and 1p (*ρ* = 0.21) models. A plot similar to Figure 5 for the 1p and 3p models is in **??** (Figure A2).

**Figure 5:**
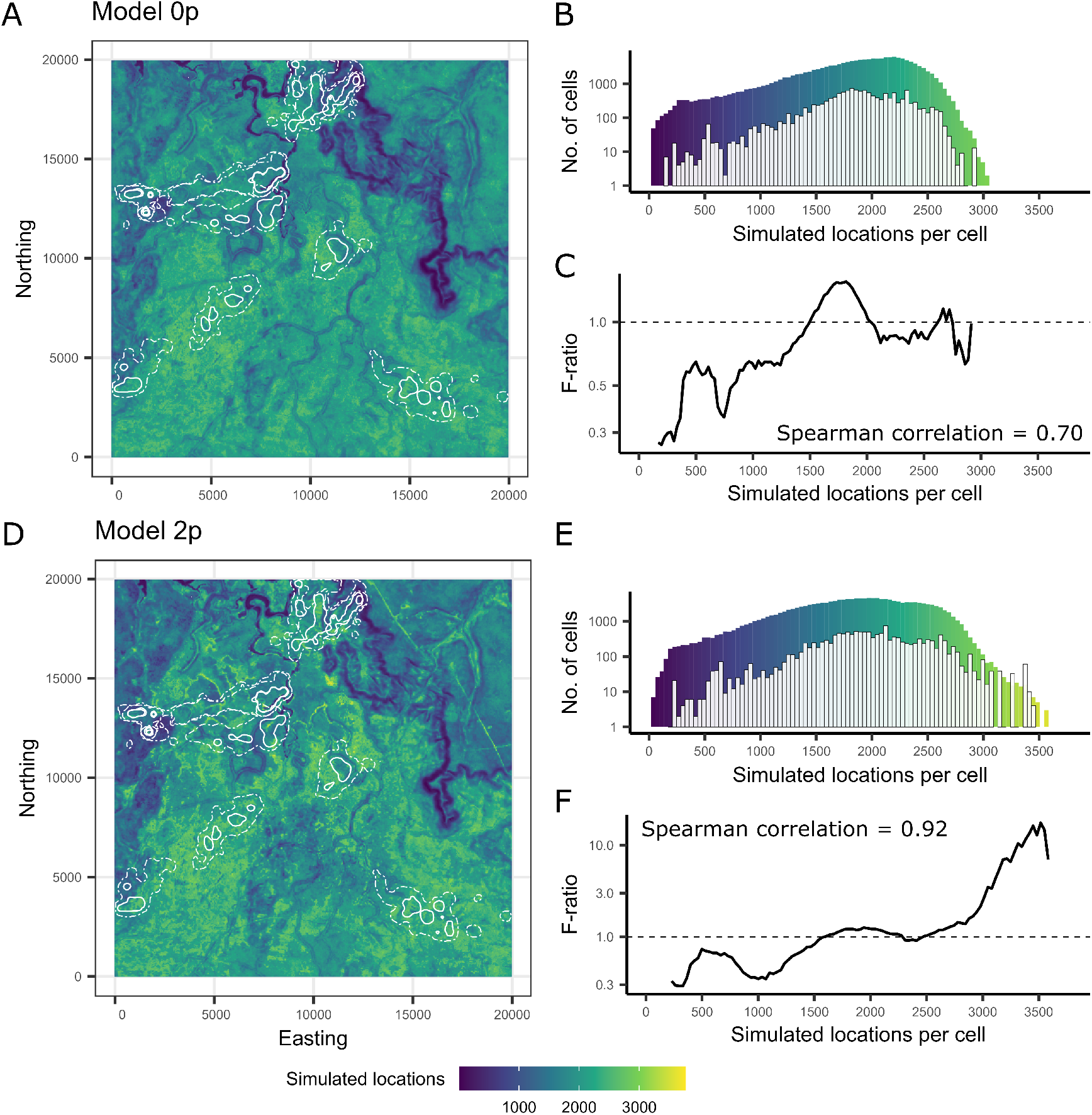
Landscape-scale predictions and their validation: Panels A and D show landscape-scale predictions for late-dry season 2018 for the static model (0p), and a dynamic model with two pairs of harmonics (2p). The density of the observed buffalo locations are shown as white contours representing the 50% (solid) and 95% (dashed) isopleths, derived using kernel density estimation fitted to each individual (n = 13,933 GPS locations in total). We considered the number of simulated locations in each cell as the landscape-scale predictions of buffalo spatial distribution. Predictions were generated by simulating 100,000 trajectories with 3,000 steps each. Panels B and E each show histograms of the predicted and observed density of locations. The coloured bars show the density of simulated locations that fell within each cell, and the white bars show the distribution of the observed buffalo locations when overlaid on these predictions. The F-ratio shown in Panels C and F is the ratio of these densities (normalised by the total number of simulated and observed locations, respectively) within a sweeping window, known as the continuous Boyce index (cBI) (Boyce et al., 2002; Hirzel et al., 2006). For a model that performs well, we would expect fewer locations in areas that are predicted to be unsuitable habitat (resulting in an F-ratio less than 1), and more observed locations in areas that are predicted to be suitable habitat (resulting in an F-ratio greater than 1). We assessed whether there was an increasing trend of the F-ratio by using the Spearman rank correlation coefficient, *ρ*, which ranges from −1 to 1, with 0 indicating a random distribution of observed locations, and a value of 1 indicating perfect predictions (denoted by a strictly monotonic increasing F-ratio). In this case the static (0p) model’s predictions were more homogeneous, likely due to averaging over the fine-scale temporal dynamics, whereas the dynamic (2p) model revealed emergent areas of high use in the landscape which may be at higher risk of environmental damage. The high-use areas in the 2p model also correlated with a high relative proportion of buffalo locations, resulting in a higher value of *ρ*.

The hourly predictions indicate that the 2p and 3p models were most accurate throughout the day, and that each model was most accurate towards the middle of the day (Figure 6), which correlates with the alignment of the mean values of the observed and simulated locations for those hours (Figure 4). The dynamic models reveal a concentration of buffalo during the middle of the day, presumably in denser canopy with high values of NDVI, which is supported by a concentration of observed buffalo locations into smaller clusters. Animations of the hourly predictions are in the online supplementary materials.

**Figure 6:**
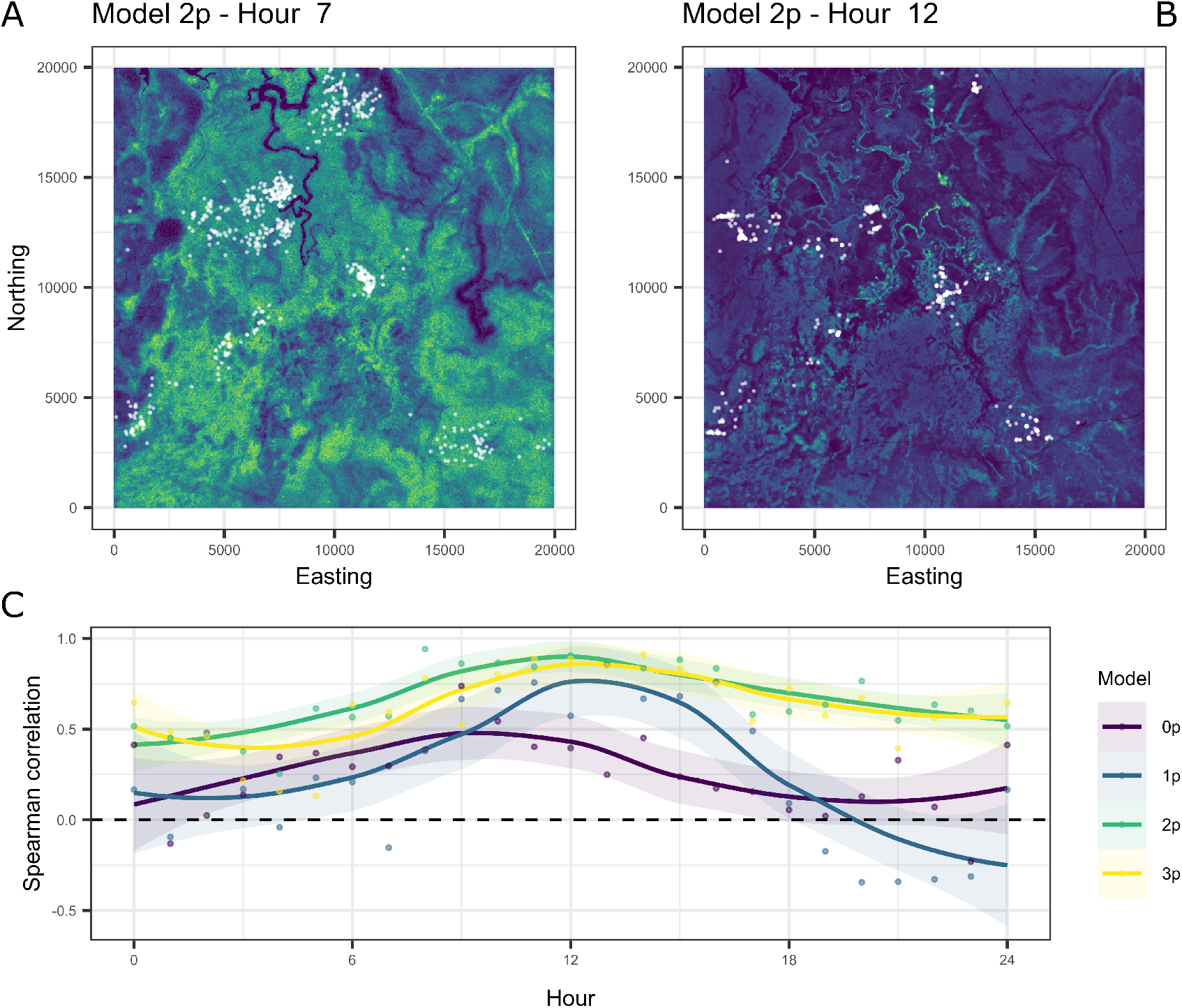
As the simulations were temporally dynamic on a daily time-scale, we generated landscape-scale predictions for each hour of the day. In Panels A and B we show landscape-scale predictions from a dynamic model with 2 pairs of harmonics (2p) for Hour 7 (7am) and midday (Hour 12). Hour 7 is during the buffalo’s period of high movement activity, which resulted in diffuse predictions that aligned with more open habitat and correlated with more spread out observed buffalo GPS locations (shown as white points, n = 657). Hour 12 is during the period when buffalo seek denser vegetation and more closed canopy to shelter from high temperatures and sun, resulting in predictions that were localised in certain areas, which correlated more closely with the tightly clustered observed buffalo GPS locations (n = 539). To aid visualisation the colours are scaled within each map, and the maximum number of simulated locations per cell are 177 and 511 for the Hour 7 and Hour 12 maps, respectively. Panel C shows the Spearman rank correlation coefficient, *ρ*, for each of the models for every hour of the day. Similar to the results shown in Figure 4, the predictions were most accurate for all models during the middle of the day, and the dynamic models with 2 and 3 pairs of harmonics (2p and 3p) were the most consistently accurate across all hours of the day. There are animations of the hourly predictions from each of the models in the online supplementary materials.

## 4 Discussion

In this study we incorporated fine-scale temporal dynamics into step selection functions (SSFs) for the purpose of generating temporally fine-scale and long-term predictions of water buffalo (*Bubalus bubalis*) distribution. We also assessed whether including fine-scale temporal dynamics allowed for more accurate long-term predictions and the identification of emergent features. The hourly summary statistics indicated that the dynamic models more realistically replicated the daily behaviours of buffalo, which resulted in more accurate landscape-scale predictions across the hours of the day, and importantly, more accurate predictions when considering longer time-scales. Our simulations also identified high-use areas in the landscape in places that may be most susceptible to environmental damage, which were not present in the static model due to an averaging of the movement and selection processes. The flexible, temporally dynamic responses in the estimated step selection parameters revealed daily patterns of buffalo movement and external selection behaviour.

Our results showed that, in the middle of the day, buffalo had strong selection for woody vegetation that is characterised by a denser canopy and high values of NDVI, which likely relates to thermoregulation. We also identified two high-movement periods near dawn and dusk, which likely describes the movement between foraging areas and the denser canopy areas for thermoregulation. For guiding management operations, our hourly predictions revealed a highly variable dynamic distribution, with diffuse predictions during the high-movement periods and concentrated use in the middle of the day. Despite the buffalo occupying a relatively small area in the middle of the day, they may be difficult to see from the air due to the dense canopy. Therefore, aerial operations such as population surveys, shooting and aggregating individuals for mustering may be more effective during the high-movement periods when buffalo are in the open, even though they may be spread over a larger area. The long-term predictions of the more accurate dynamic models showed there are several areas that may be at higher risk of damage (identified by high predicted use). If these areas are ecologically or culturally important, then persistent management actions such as exclusion fencing may be considered (Ens et al., 2016; Sloane et al., 2024).

Given the pervasiveness of fine-scale temporal dynamics in animal behaviour, our results show that we may be missing out on valuable information when generating predictions. These findings may be even more pertinent to species that have greater mobility (e.g. flighted animals), central place foragers, or for other species that may use distinctly different areas throughout the course of a day (Chapman et al., 1989; Leblond et al., 2010; Kohl et al., 2018; Ylitalo et al., 2021). In cases where temporally fine-scale predictions might be useful, such as predicting human-wildlife interactions (Carter et al., 2012; Buderman et al., 2018), identifying areas of possible zoonotic disease transmission (Parsons et al., 2014), mitigating fishing impacts (Ferńandez & Anderson, 2000; Ouled-Cheikh et al., 2020), or intensive conservation management operations, we have shown that incorporating temporal dynamics into simulated trajectories can be a valuable approach to meet these objectives. Including the temporally dynamic processes into the model may also result in identifying emergent features of the aggregated behaviour of the population, even when the temporally fine-scale predictions are not of interest. We also expect that fine-scale temporal dynamics may affect other prediction targets, such as connectivity and movement corridors, particularly due to dynamic representation of movement behaviour (Hooker et al., 2021; Whittington et al., 2022; Aiello et al., 2023; Hofmann et al., 2023; Sells et al., 2023).

Given the potential influence on predictions, assessing how temporally dynamic the movement and habitat selection behaviour is would be a useful exploratory step prior to model fitting. It is possible to identify the trend of temporal dynamics through the dynamic summary statistic approach we took for the step lengths and habitat covariates for each hour (Figure 4). In our case, the estimated coefficients of NDVI, canopy cover and herbaceous vegetation exhibited a single mode of selection, and all dynamic models (denoted 1p, 2p and 3p for the number of harmonic pairs) largely replicated this. However, for the movement behaviour, there was a multimodal pattern that was only captured by the more flexible models that allowed for multiple modes (2p and 3p), likely contributing to the higher prediction accuracy for these models.

In the models presented in this paper we used harmonic terms to incorporate the temporally dynamic behaviour into the SSFs, although a recent paper by Klappstein et al. (2024) shows that splines are another natural option, which also allow for SSF extensions and hierarchical model fitting using the mgcv package (Wood, 2011). Simulating trajectories from dynamic models fitted with spline terms would be the same process as we have presented here, and would just be a matter of indexing the coefficients for *τ* ∈ *T* when simulating trajectories. It would also be possible to fit the two-dimensional selection surfaces that we achieved by interacting quadratic terms and harmonics by using interacting spline terms, which would be more flexible than our approach, although it would add more parameters into the model. An additional benefit of the spline approach is that the temporal dynamics do not need to be cyclic, which can allow for temporally dynamic movement and habitat selection to be inferred and simulated from in cases such as post-release behaviour and home range establishment (Maor-Cohen et al., 2021; Picardi et al., 2021; Cisneros-Araujo et al., 2024).

Another alternative method of incorporating temporal dynamics into animal movement models is through state-switching models such as a hidden Markov model (HMM) (Langrock et al., 2012; McClintock et al., 2012). In HMMs, behaviours such as foraging, resting and transiting are represented as states with different movement parameters, and when combined with SSFs (HMM-SSF/HMM-iSSA), different habitat selection parameters (Picardi et al., 2022; Beumer et al., 2023; Klappstein et al., 2023; Pohle et al., 2024). These models can easily incorporate temporal dynamics as the transition matrix governing state-switching can depend on time, although in HMMs the states are discrete, whereas real behaviour changes may be gradual and continuous, which may affect predictions in some cases. However, hierarchical HMMs may be an effective method to incorporate behavioural changes over multiple time-scales, and when combined with habitat selection, may produce informative and accurate simulation models (Leos-Barajas et al., 2017; Adam et al., 2019).

Simulated trajectories of animal movement have applications beyond what is possible compared to analytic prediction approaches such as resource selection functions (Potts & Börger, 2023), and we see many potential applications of using SSFs to generate trajectories, particularly when they can capture realistic behavioural dynamics. A promising application is for counter-factual analysis, where landscape covariates can be modified to represent a disturbance, management action, reserve or habitat corridor design, and trajectories can be simulated to understand how animals may respond. Other applications include near-term trajectories that assess the likelihood of an area being used by an animal in the future, which may be useful to assess colonisation and invasion potential of introduced species (Lustig et al., 2017, 2019; Patterson et al., 2024), or to plan upcoming management when the locations of animals are currently known. Trajectories from SSFs have already seen useful application for connectivity, and there is potential to identify movement corridors and understand metapopulation dynamics in fragmented environments (Hooker et al., 2021; Whittington et al., 2022; Aiello et al., 2023; Hofmann et al., 2023; Sells et al., 2023), particularly when combined with additional data such as genetics to assess historic connectivity (Lowe & Allendorf, 2010; Dussex et al., 2015).

An additional feature of parameterised SSFs is that it can serve as the foundation for a more complex model (Hauenstein et al., 2019). Additional parameters that govern behavioural and ecological factors such as more sophisticated memory effects and home range behaviour, social interactions, and more flexible relationships between the animal and resource covariates can be added for a more sophisticated simulation model. The model would then need to be calibrated to data, but the SSF estimates for the movement and resource selection parameters can be used as priors, and approaches such as simulation-based inference can be used to fit the updated model to data (Hartig et al., 2011; Cranmer et al., 2020; Tejero-Cantero et al., 2020). For an example of this approach see Hauenstein et al. (2019).

## 5 Conclusions

We have generated temporally fine-scale predictions of animal spatial distribution via simulations from step selection models that incorporated fine-scale temporal dynamics through harmonic terms. We found that simulations from dynamic models replicated the observed behaviour more closely, which resulted in more accurate temporally fine-scale and long-term predictions of distribution. Adding temporal dynamics to the movement behaviour and memory processes also allowed for useful inference towards our study species, the invasive water buffalo (*Bubalus bubalis*), which can help to guide management operations in Northern Australia. As more ecologists are turning to SSFs to simulate trajectories for understanding future distribution and connectivity, we suggest incorporating temporal dynamics to more realistically represent animal behaviour through time, which can improve predictions and identify emergent features that would be otherwise missed. For those wanting to better understand movement and habitat selection of their species, including temporal dynamics can provide richer information, particularly for species with clear daily or seasonal patterns of behaviour.

## Acknowledgements

SWF was supported by an Australian Government Research Training Program Scholarship and a CSIRO top-up scholarship. CD was supported by an Australian Research Council Future Fellowship (FT210100260). This work was funded by the federal government’s Department of Agriculture, Fisheries and Forestry under the Control tools and technologies for established pest animals and weeds competitive grants program and the Smart Farming Partnerships Program (round 2). Computational resources and services used in this work were provided by the eResearch Office, Queensland University of Technology, Brisbane, Australia. Landsat-7 image courtesy of the U.S. Geological Survey. We thank Michael Lynch (Melbourne Zoo), Jack Larson and Patrick Carmody for their assistance in the capture and collaring of animals. The Ricky Archer and the Bawinanga Rangers for their support in all field activities. SWF thanks Charlotte Patterson and members of the Applied Mathematical Ecology (AMEG) and Bayesian Research and Applications Group (BRAG) for helpful discussions and feedback on the work.

## Author Contributions

SWF, DP, JRP and AJH conceived the ideas and developed the modelling methodology; AJH, JP and EV designed the data collection and collected the data; SF analysed the data and led the writing of the manuscript. All authors contributed critically to the drafts and gave final approval for publication.

## Data Availability Statement

Code and data are available at https://github.com/swforrest/dynamic SSF sims, including a walk-through of fitting harmonic terms in step selection models. For peer-review and access to larger files such as spatial covariates and intermediate outputs a zipped folder has been uploaded to Zenodo. The URL to access the restricted Zenodo folder is: https://zenodo.org/records/10838069.

## Appendix A Additional results and simulation convergence

**Figure A1:**
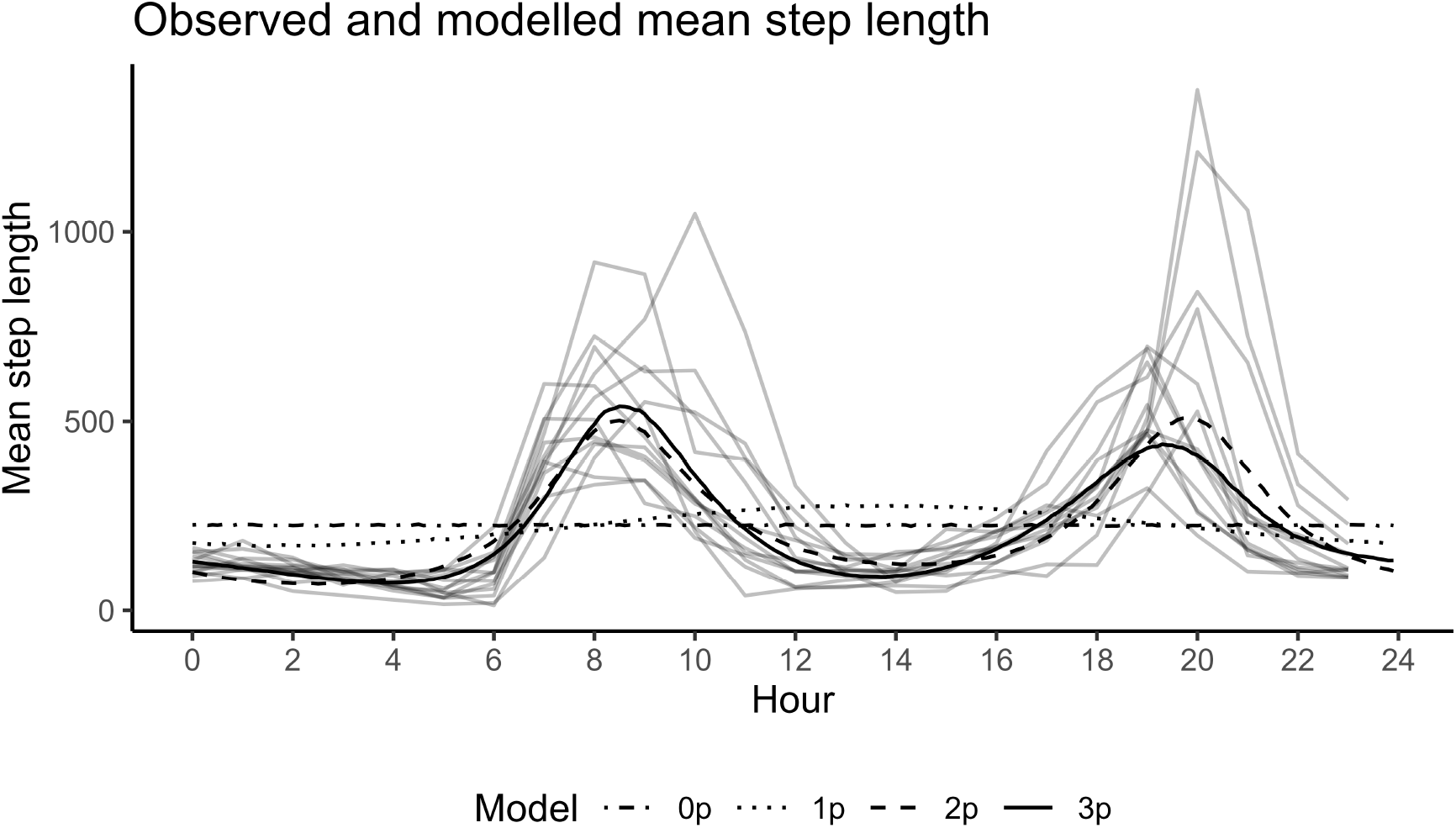
The observed and modelled mean step lengths across the hours of the day from all models when sampling from the estimated movement kernels. The observed mean step lengths are shown as a grey line for each individual. Samples from the estimated, temporally dynamic, step length distribution are shown with the black lines, with 0p, 1p, 2p and 3p referring to the number of pairs of harmonic terms included in the model. The movement parameters were updated using the estimated coefficients and the tentative gamma and von Mises distributions that were fitted to the observed data and that random steps were sampled from prior to model fitting (Avgar et al., 2016; Fieberg et al., 2021; Michelot et al., 2024). The model with a single pair of harmonics is restricted to a single period, and fails to capture the two peaks of movement in the observed data. The 2p and 3p models are similar, although the model with three harmonics has more flexibility to represent the difference between the dawn and dusk periods of high activity.

**Figure A2:**
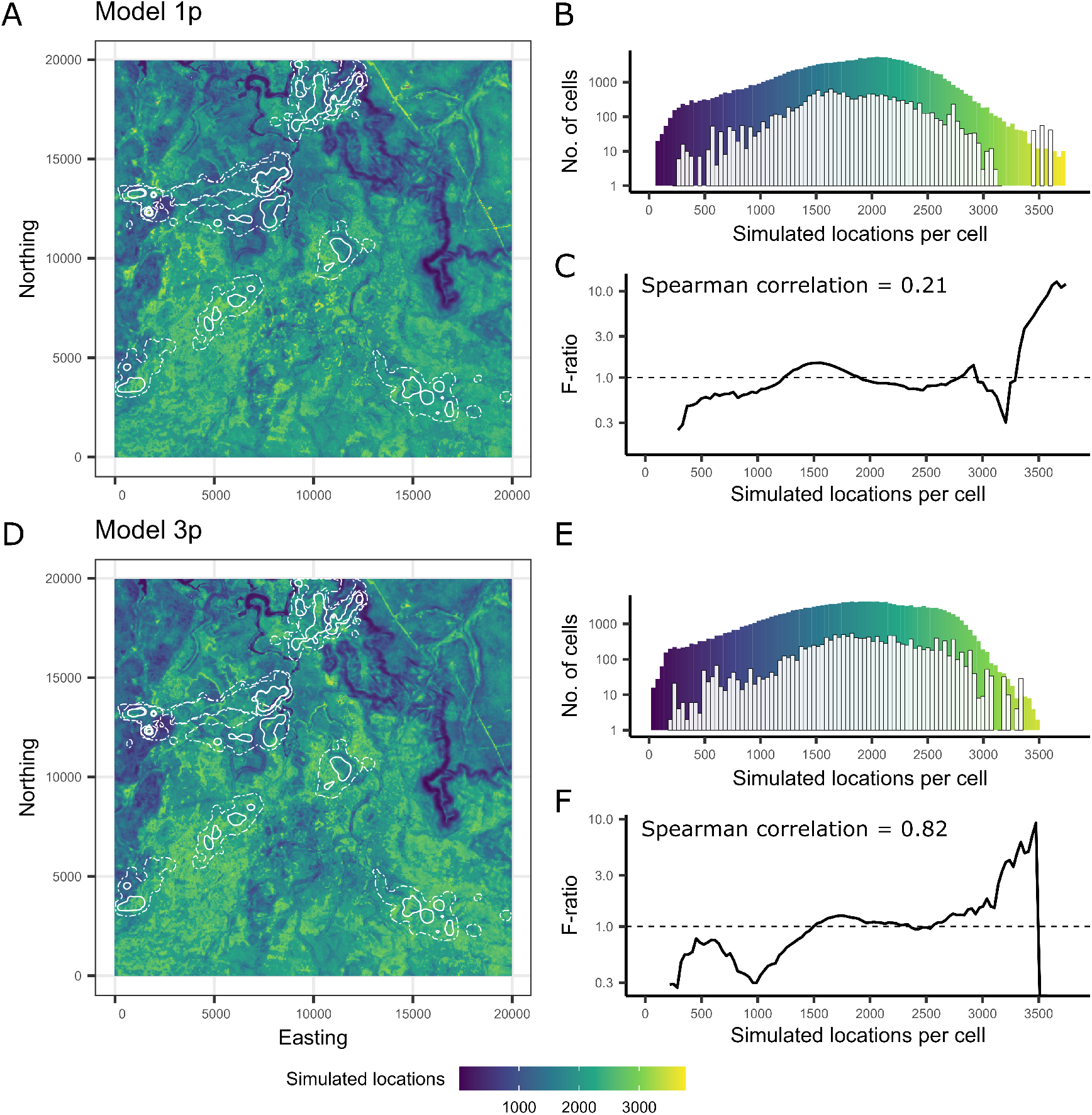
Landscape-scale predictions and their validation: Panels A and D show landscape-scale predictions for late-dry season 2018 for the model with 1 pair of harmonics (1p), and a dynamic model with 3 pairs of harmonics (3p). The density of the observed buffalo locations are shown as white contours representing the 50% (solid) and 95% (dashed) isopleths, derived using kernel density estimation fitted to each individual (n = 13,933 GPS locations in total). We considered the number of simulated locations in each cell as the landscape-scale predictions of buffalo spatial distribution. Predictions were generated by simulating 100,000 trajectories with 3,000 steps each. Panels B and E each show histograms of the predicted and observed density of locations. The coloured bars show the density of simulated locations that fell within each cell, and the white bars show the distribution of the observed buffalo locations when overlaid on these predictions. The F-ratio shown in Panels C and F is the ratio of these densities (normalised by the total number of simulated and observed locations, respectively) within a sweeping window, known as the continuous Boyce index (cBI) (Boyce et al., 2002; Hirzel et al., 2006). For a model that performs well, we would expect fewer locations in areas that are predicted to be unsuitable habitat (resulting in an F-ratio less than 1), and more observed locations in areas that are predicted to be suitable habitat (resulting in an F-ratio greater than 1). We assessed whether there was an increasing trend of the F-ratio by using the Spearman rank correlation coefficient, *ρ*, which ranges from −1 to 1, with 0 indicating a random distribution of observed locations, and a value of 1 indicating perfect predictions (denoted by a strictly monotonic increasing F-ratio). In this case the static (0p) model’s predictions were more homogeneous, likely due to averaging over the fine-scale temporal dynamics, whereas the dynamic (2p) model revealed emergent areas of high use in the landscape which may be at higher risk of environmental damage. The high-use areas in the 2p model also correlated with a high relative proportion of buffalo locations, resulting in a higher value of *ρ*.

**Figure A3:**
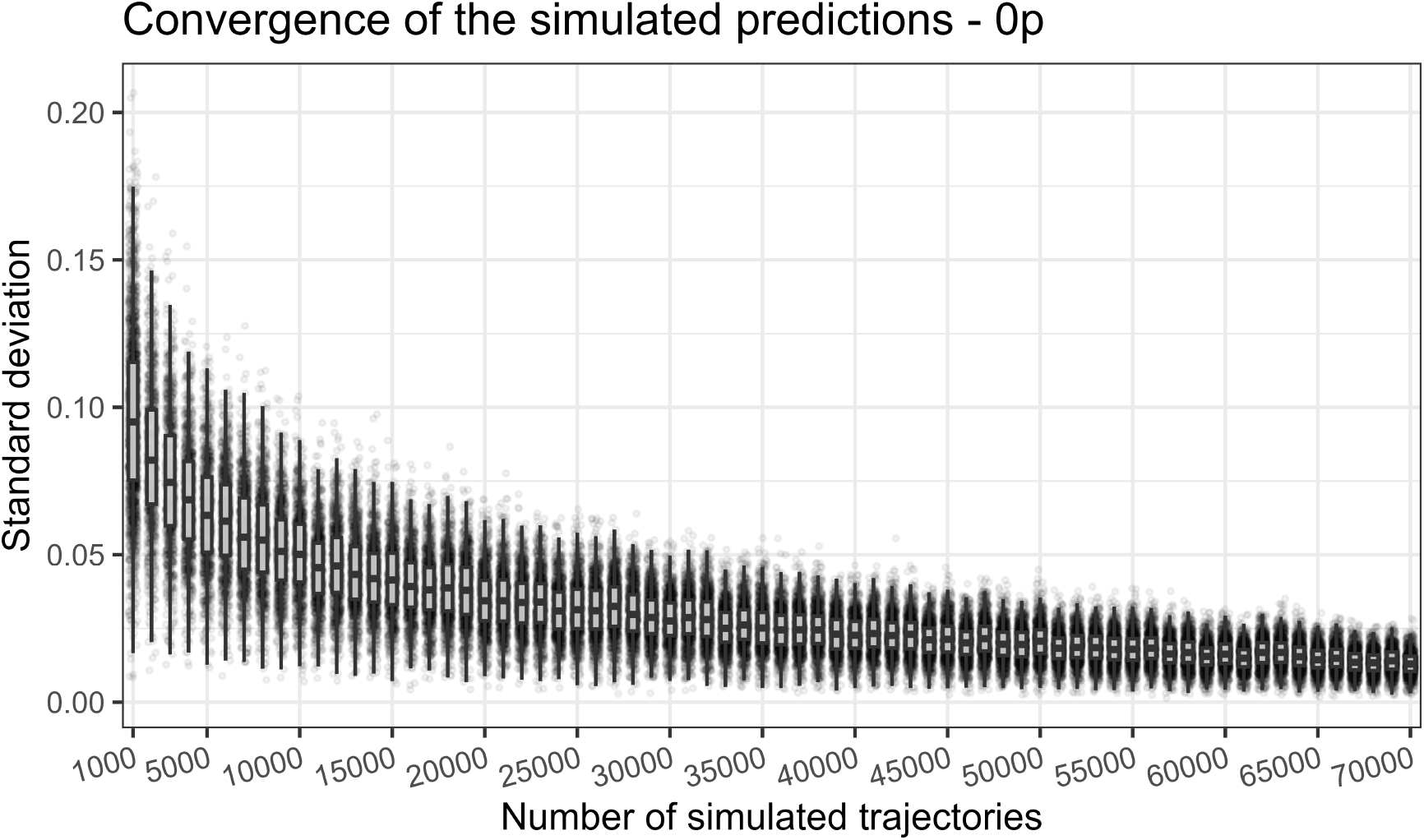
We assessed the convergence of the simulations by randomly sampling 10 subsets of increasing numbers of trajectories (without replacement), and calculating the standard deviation of normalised prediction values (from 0 to 1) in 1000 randomly distributed cells. Here each point represents the standard deviation across the 10 subsets for 1 of the randomly distributed cells. This plot shows the convergence of the 2p simulations, although the convergence was similar for all models. It should be noted that as the number of trajectories in each subset approaches the size of the full dataset (100,000 trajectories), the subsets will more closely represent the full dataset, reducing the variance between subsets.

## Appendix B Considerations when simulating animal movement trajectories

It is important to note that simulation performance is sensitive to the form of the covariates. Here we included covariates that mostly range between −1 and 1 or between 0 and 1, and most of the environmental space was visited, limiting out of sample prediction. Covariates such as ‘distance to’ covariates can produce different behaviour in simulations, as distances can become very large, leading to very high or low selection behaviour (depending on the direction of the coefficient) which only becomes apparent when simulating trajectories or predicting with other approaches. In these cases we suggest considering transformations of the covariate, such as log-transformation or similar. It should be noted however, that ‘distance to’ covariates can be well-suited to temporally dynamic simulations as they can replicate behaviour such as central place foraging for instance. This would be achieved by using a ‘distance to a central place(s)’ covariate, and the temporally dynamic coefficients would allow for there to be avoidance (selecting for larger distances to the central place) when setting out for foraging for instance, which becomes attraction as the coefficient crosses 0, allowing for the simulated individuals to return to their central place(s).

We also recommend quadratic terms in the models used for simulations, as it allows for the simulated individuals to select within a region of the covariates, rather than monotonically increasing or decreasing values as is the case when only considering linear terms. An example from our case is for NDVI (Figure 3), where even for the static (0p) model, selection is highest for NDVI values between 0.25 and 0.5, and there is avoidance for values less than 0 and greater than 0.7. Assessing the accuracy of simulations to different covariates is therefore important, and we suggest following a summary statistic approach as we have taken here, giving more weight to summary statistics that represent movement characteristics relating to the research question. An approach to assess the more qualitative aspects of the overall trajectory would be the ‘lineup’ approach (Fieberg et al., 2023), as many of the trajectories appeared visually similar to the observed data. We also advocate for using simulations as a means of model selection and to investigate drivers of animal movement decisions (Potts et al., 2022).

In our case we did not consider parameter uncertainty of the model outputs in our simulations. This would be computationally expensive (as many more simulations would be required), although it would be straightforward to achieve, particularly if using a Bayesian sampling approach such as Markov Chain Monte Carlo (MCMC) to fit the models. In that case, samples could be drawn from the joint posterior and passed to the simulation procedure, essentially resulting in posterior predictions of animal movement trajectories (McElreath, 2020). For studies where predictions are generated to assess a research question such as distribution or connectivity, incorporating this uncertainty may be important, particularly if models are fitted hierarchically and one wants to consider the individual variability of spatial behaviours (Dall et al., 2012; Bastille-Rousseau & Wittemyer, 2019, 2022; Stuber et al., 2022). An alternative approach to consider individual variability is to fit models to each individual animal separately, which can each be used to simulate many trajectories. However in our case, fitting models to individuals separately led to pathological behaviour in the movement parameter updating process, which resulted in impossible movement kernels in some cases. For a more in-depth discussion see Appendix C.

## Appendix C Issues when updating temporally dynamic movement parameters

When fitting models to individuals separately, we had issues for some individuals where the movement parameter ‘updating’ (Avgar et al., 2016; Fieberg et al., 2021) resulted in gamma distributions that had impossibly large step lengths. Using the inferred parameters of the step selection model, the updated gamma scale parameter of the model, *q*^, which is defined as 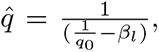 is the tentative scale parameter and *β_l_* is the coefficient of the step length covariate, approached 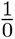, resulting in extremely large scale parameters. We also noticed that the scale parameter was accounting for the high variation in step lengths throughout the day, and the shape parameter remained relatively constant. The stability of the shape parameter suggests that the scale parameter is ‘absorbing’ the temporal variation, resulting in pathological results, although we also observed similar behaviour when using the exponential distribution, where only a single parameter is estimated. It is likely that further increasing the number of random steps will increase the accuracy of the numerical integration (Michelot et al., 2024), although even with 100 random steps we observed the pathological behaviour. For a high number of parameters and a large dataset increasing the number of steps beyond this can be computationally prohibitive, particularly when fitting models hierarchically. When using the TwoStep estimation for all individuals this was not an issue however, as it was only the model fits for some of the individuals when fitted separately that resulted in the pathological behaviour.

It is clear that this behaviour is due to the large difference between the ‘tentative’ step length and those that are estimated from the model due to the significant temporal changes. For other species that have less dynamic movement changes when compared to the buffalo this isn’t likely to be an issue. The behaviour may also indicate an issue with harmonic terms, which are constrained by their sinusoidal behaviour, and may ‘overshoot’ at the modes (and approach 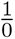) in an attempt to accommodate sharp changes in movement behaviour, and either more harmonics that lend more flexibility, or an approach such as splines may lead to better results in these cases.

## References

Adam, T., Griffiths, C. A., Leos-Barajas, V., Meese, E. N., Lowe, C. G., Blackwell, P. G., Righton, D., & Langrock, R. (2019). Joint modelling of multi-scale animal movement data using hierarchical hidden markov models. Methods in Ecology and Evolution, 10 (9), 1536–1550. 10.1111/2041-210x.13241

Ager, A. A., Johnson, B. K., Kern, J. W., & Kie, J. G. (2003). Daily and seasonal movements and habitat use by female rocky mountain elk and mule deer. Journal of Mammalogy, 84 (3), 1076–1088. 10.1644/BBa-020

Aiello, C. M., Galloway, N. L., Prentice, P. R., Darby, N. W., Hughson, D., & Epps, C. W. (2023). Movement models and simulation reveal highway impacts and mitigation opportunities for a metapopulation-distributed species. Landscape Ecology, 38 (4), 1085–1103. 10.1007/s10980-023-01600-6

Avgar, T., Potts, J. R., Lewis, M. A., & Boyce, M. (2016). Integrated step selection analysis: Bridging the gap between resource selection and animal movement. Methods in Ecology and Evolution, 7 (5), 619–630. 10.1111/2041-210x.12528

Beumer, L. T., Schmidt, N. M., Pohle, J., Signer, J., Chimienti, M., Desforges, J.-P., Hansen, L. H., Højlund Pedersen, S., Rudd, D. A., Stelvig, M., & van Beest, F. M. (2023). Accounting for behaviour in fine-scale habitat selection: A case study highlighting methodological intricacies. The Journal of Animal Ecology, 92 (10), 1937–1953. 10.1111/1365-2656.13984

Boyce, M., Pitt, J., Northrup, J. M., Morehouse, A. T., Knopff, K. H., Cristescu, B., & Stenhouse, G. B. (2010). Temporal autocorrelation functions for movement rates from global positioning system radiotelemetry data. Philosophical Transactions of the Royal Society of London. Series B, Biological Sciences, 365 (1550), 2213–2219. 10.1098/rstb.2010.0080

Boyce, M., Vernier, P. R., Nielsen, S. E., & Schmiegelow, F. K. A. (2002). Evaluating resource selection functions. Ecological Modelling, 157 (2), 281–300. 10.1016/S0304-3800(02)00200-4

Brooks, M. E., Kristensen, K., van Benthem, K. J., Magnusson, A., Berg, C. W., Nielsen, A., Skaug, H. J., Mächler, M., & Bolker, B. M. (2017). glmmTMB balances speed and flexibility among packages for zero-inflated generalized linear mixed modeling. The R Journal, 9 (2), 378–400.

Buderman, F. E., Hooten, M. B., Alldredge, M. W., Hanks, E. M., & Ivan, J. S. (2018). Time-varying predatory behavior is primary predictor of fine-scale movement of wildland-urban cougars. Movement Ecology, 6, 22. 10.1186/s40462-018-0140-6

Campbell, H. A., Loewensteiner, D. A., Murphy, B. P., Pittard, S., & McMahon, C. R. (2020). Seasonal movements and site utilisation by asian water buffalo (bubalus bubalis) in tropical savannas and floodplains of northern australia. Wildlife Research, 48 (3), 230–239. 10.1071/WR20070

Carter, N. H., Shrestha, B. K., Karki, J. B., Pradhan, N. M. B., & Liu, J. (2012). Coexistence between wildlife and humans at fine spatial scales. Proceedings of the National Academy of Sciences of the United States of America, 109 (38), 15360–15365. 10.1073/pnas.1210490109

Chapman, C. A., Chapman, L. J., & McLaughlin, R. L. (1989). Multiple central place foraging by spider monkeys: Travel consequences of using many sleeping sites. Oecologia, 79 (4), 506–511. 10.1007/BF00378668

Cisneros-Araujo, P., Garrote, G., Corradini, A., Farhadinia, M. S., Robira, B., López, G., Fernández, L., López-Parra, M., García-Tardío, M., Arenas-Rojas, R., del Rey, T., Salcedo, J., Sarmento, P., Sánchez, J. F., Palacios, M. J., García-Viñás, J. I., Damiani, M. L., Hachem, F., Gastón, A., & Cagnacci, F. (2024). Born to be wild: Captive-born and wild iberian lynx (lynx pardinus) reveal space-use similarities when reintroduced for species conservation concerns. Biological Conservation, 294, 110646. 10.1016/j.biocon.2024.110646

Craiu, R. V., Duchesne, T., Fortin, D., & Baillargeon, S. (2011). Conditional logistic regression with longitudinal follow-up and Individual-Level random coefficients: A stable and efficient Two-Step estimation method. Journal of Computational and Graphical Statistics, 20 (3), 767–784. 10.1198/jcgs.2011.09189

Craiu, R. V., Duchesne, T., Fortin, D., & Baillargeon, S. (2016). R package ‘TwoStepCLogit’. conditional logistic regression: A Two-Step estimation method (tech. rep.).

Cranmer, K., Brehmer, J., & Louppe, G. (2020). The frontier of simulation-based inference. Proceedings of the National Academy of Sciences of the United States of America, 117 (48), 30055–30062. 10.1073/pnas.1912789117

Currey, L. M., Heupel, M. R., Simpfendorfer, C. A., & Williams, A. J. (2015). Assessing fine-scale diel movement patterns of an exploited coral reef fish. Animal Biotelemetry, 3 (1), 41. 10.1186/s40317-015-0072-5

Dussex, N., Sainsbury, J., Moorhouse, R., Jamieson, I. G., & Robertson, B. C. (2015). Evidence for bergmann’s rule and not allopatric subspeciation in the threatened kaka (nestor meridionalis). The Journal of Heredity, 106 (6), 679–691. 10.1093/jhered/esv079

Eisaguirre, J. M., Williams, P. J., & Hooten, M. B. (2024). Rayleigh step-selection functions and connections to continuous-time mechanistic movement models. Movement Ecology, 12 (1), 14. 10.1186/s40462-023-00442-w

Ellison, N., Potts, J. R., Strickland, B. K., Demarais, S., & Street, G. M. (2024). Combining animal interactions and habitat selection into models of space use: A case study with white-tailed deer. *Wildlife Biology*, e01211. 10.1002/wlb3.01211

Ens, E. J., Daniels, C., Nelson, E., Roy, J., & Dixon, P. (2016). Creating multi-functional land-scapees: Using exclusion fences to frame feral ungulate management preferences in remote aboriginal-owned northern australia. Biological Conservation, 197, 235–246. 10.1016/j.biocon.2016.03.007

Fagan, W. F., Lewis, M. A., Auger-Méthé, M., Avgar, T., Benhamou, S., Breed, G., LaDage, L., Schlägel, U. E., Tang, W.-W., Papastamatiou, Y. P., Forester, J., & Mueller, T. (2013). Spatial memory and animal movement. Ecology Letters, 16 (10), 1316–1329. 10.1111/ele.12165

Fernández, P., & Anderson, D. J. (2000). Nocturnal and diurnal foraging activity of hawaiian albatrosses detected with a new immersion monitor. The Condor, 102 (3), 577–584. 10.1093/condor/102.3.577

Fieberg, J., Signer, J., Smith, B., & Avgar, T. (2021). A ‘how to’ guide for interpreting parameters in habitat-selection analyses. The Journal of Animal Ecology, 90 (5), 1027–1043. 10.1111/1365-2656.13441

Forester, J. D., Im, H. K., & Rathouz, P. J. (2009). Accounting for animal movement in estimation of resource selection functions: Sampling and data analysis. Ecology, 90 (12), 3554–3565. 10.1890/08-0874.1

Forrest, S. W., Rodŕıguez-Recio, M., & Seddon, P. J. (2024). Home range and dynamic space use reveals age-related differences in risk exposure for reintroduced parrots. *Conservation Science and Practice*, e13119. 10.1111/csp2.13119

Fortin, D., Beyer, H. L., Boyce, M. S., Smith, D. W., Duchesne, T., & Mao, J. S. (2005). Wolves influence elk movements: Behavior shapes a trophic cascade in yellowstone national park. Ecology, 86 (5), 1320–1330. 10.1890/04-0953

Fox, R. J., & Bellwood, D. R. (2011). Unconstrained by the clock? plasticity of diel activity rhythm in a tropical reef fish, siganus lineatus. Functional Ecology, 25 (5), 1096–1105. 10.1111/j.1365-2435.2011.01874.x

Fryxell, J. M., Hazell, M., Börger, L., Dalziel, B. D., Haydon, D. T., Morales, J. M., McIntosh, T., & Rosatte, R. C. (2008). Multiple movement modes by large herbivores at multiple spatiotemporal scales. Proceedings of the National Academy of Sciences of the United States of America, 105 (49), 19114–19119. 10.1073/pnas.0801737105

Hanks, E. M., Hooten, M. B., & Alldredge, M. W. (2015). Continuous-time discrete-space models for animal movement. The Annals of Applied Statistics, 9 (1), 145–165. 10.1214/14-AOAS803

Hartig, F., Calabrese, J. M., Reineking, B., Wiegand, T., & Huth, A. (2011). Statistical inference for stochastic simulation models–theory and application. Ecology Letters, 14 (8), 816–827. 10.1111/j.1461-0248.2011.01640.x

Hauenstein, S., Fattebert, J., Grüebler, M. U., Naef-Daenzer, B., Pe’er, G., & Hartig, F. (2019). Calibrating an individual-based movement model to predict functional connectivity for little owls. Ecological Applications, 29 (4), e01873. 10.1002/eap.1873

Hijmans, R. J. (2024). Spatial data analysis [r package terra version 1.7–71].

Hirzel, A. H., Le Lay, G., Helfer, V., Randin, C., & Guisan, A. (2006). Evaluating the ability of habitat suitability models to predict species presences. Ecological Modelling, 199 (2), 142– 152. 10.1016/j.ecolmodel.2006.05.017

Hofmann, D. D., Cozzi, G., McNutt, J. W., Ozgul, A., & Behr, D. M. (2023). A three-step approach for assessing landscape connectivity via simulated dispersal: African wild dog case study. Landscape Ecology, 38 (4), 981–998. 10.1007/s10980-023-01602-4

Hooker, M. J., Clark, J., Bond, B. T., & Chamberlain, M. J. (2021). Evaluation of connectivity among american black bear populations in georgia. The Journal of Wildlife Management, 85 (5), 979–988. 10.1002/jwmg.22041

Horn, B. K. P. (1981). Hill shading and the reflectance map. Proceedings of the IEEE, 69 (1), 14–47. 10.1109/PROC.1981.11918

Klappstein, N. J., Thomas, L., & Michelot, T. (2023). Flexible hidden markov models for behaviour-dependent habitat selection. Movement Ecology, 11 (1), 30. 10.1186/s40462-023-00392-3

Klappstein, N., Michelot, T., Fieberg, J., Pedersen, E., Field, C., & Flemming, J. M. (2024, January). Step selection analysis with non-linear and random effects in mgcv. 10.1101/2024.01.05.574363

Klappstein, N. J., Potts, J. R., Michelot, T., Börger, L., Pilfold, N. W., Lewis, M. A., & Derocher, A. E. (2022). Energy-based step selection analysis: Modelling the energetic drivers of animal movement and habitat use. The Journal of Animal Ecology, 91 (5), 946–957. 10.1111/1365-2656.13687

Kohl, M. T., Stahler, D. R., Metz, M. C., Forester, J. D., Kauffman, M. J., Varley, N., White, P. J., Smith, D. W., & MacNulty, D. R. (2018). Diel predator activity drives a dynamic landscape of fear. Ecological Monographs, 88 (4), 638–652. 10.1002/ecm.1313

Langrock, R., King, R., Matthiopoulos, J., Thomas, L., Fortin, D., & Morales, J. M. (2012). Flexible and practical modeling of animal telemetry data: Hidden markov models and extensions. Ecology, 93 (11), 2336–2342. 10.1890/11-2241.1

Leblond, M., Dussault, C., & Ouellet, J.-P. (2010). What drives fine-scale movements of large herbivores? a case study using moose. Ecography, 33 (6), 1102–1112. 10.1111/j.1600-0587.2009.06104.x

Leos-Barajas, V., Gangloff, E. J., Adam, T., Langrock, R., van Beest, F. M., Nabe-Nielsen, J., & Morales, J. M. (2017). Multi-scale modeling of animal movement and general behavior data using hidden markov models with hierarchical structures. Journal of Agricultural, Biological, and Environmental Statistics, 22 (3), 232–248. 10.1007/S13253-017-0282-9/FIGURES/5

Levin, S. A. (1992). The problem of pattern and scale in ecology: The robert h. macarthur award lecture. Ecology, 73 (6), 1943–1967. 10.2307/1941447

Lowe, W. H., & Allendorf, F. W. (2010). What can genetics tell us about population connectivity? Molecular Ecology, 19 (15), 3038–3051. 10.1111/j.1365-294X.2010.04688.x

Lustig, A., James, A., Anderson, D., & Plank, M. (2019). Pest control at a regional scale: Identifying key criteria using a spatially explicit, agent-based model. The Journal of Applied Ecology, 56 (7), 1515–1527. 10.1111/1365-2664.13387

Lustig, A., Worner, S. P., Pitt, J. P. W., Doscher, C., Stouffer, D. B., & Senay, S. D. (2017). A modeling framework for the establishment and spread of invasive species in heterogeneous environments. Ecology and Evolution, 7 (20), 8338–8348. 10.1002/ece3.2915

Maor-Cohen, M., Bar-David, S., Dolev, A., Berger-Tal, O., Saltz, D., & Spiegel, O. (2021). Settling in: Reintroduced persian fallow deer adjust the borders and habitats of their Home-Range during the first 5 years post release. Frontiers in Conservation Science, 2. 10.3389/fcosc.2021.733703

McClintock, B. T., King, R., Thomas, L., Matthiopoulos, J., McConnell, B. J., & Morales, J. M. (2012). A general discrete-time modeling framework for animal movement using multistate random walks. Ecological Monographs, 82 (3), 335–349. 10.1890/11-0326.1

McMahon, C. R., & Bradshaw, C. J. A. (2008). To catch a buffalo: Field immobilisation of asian swamp buffalo using etorphine and xylazine. Australian Veterinary Journal, 86 (6), 235– 241. 10.1111/j.1751-0813.2008.00303.x

Meese, E. N., & Lowe, C. G. (2020). Active acoustic telemetry tracking and tri-axial accelerometers reveal fine-scale movement strategies of a non-obligate ram ventilator. Movement Ecology, 8, 8. 10.1186/s40462-020-0191-3

Michelot, T., Klappstein, N. J., Potts, J. R., & Fieberg, J. (2024). Understanding step selection analysis through numerical integration. Methods in Ecology and Evolution, 15 (1), 24–35. 10.1111/2041-210x.14248

Mihailou, H., & Massaro, M. (2021). An overview of the impacts of feral cattle, water buffalo and pigs on the savannas, wetlands and biota of northern australia. Austral Ecology, 46 (5), 699–712. 10.1111/aec.13046

Morales, J. M., Moorcroft, P. R., Matthiopoulos, J., Frair, J. L., Kie, J. G., Powell, R. A., Merrill, E. H., & Haydon, D. T. (2010). Building the bridge between animal movement and population dynamics. Philosophical transactions of the Royal Society of London. Series B, Biological sciences, 365 (1550), 2289–2301. 10.1098/rstb.2010.0082

Muff, S., Signer, J., & Fieberg, J. (2020). Accounting for individual-specific variation in habitat-selection studies: Efficient estimation of mixed-effects models using bayesian or frequentist computation (E. V. Wal, Ed.). The Journal of Animal Ecology, 89 (1), 80–92. 10.1111/1365-2656.13087

Munden, R., Börger, L., Wilson, R. P., Redcliffe, J., Brown, R., Garel, M., & Potts, J. R. (2021). Why did the animal turn? time-varying step selection analysis for inference between observed turning points in high frequency data. Methods in Ecology and Evolution, (2041–210X.13574). 10.1111/2041-210x.13574

Myneni, R. B., Hall, F. G., Sellers, P. J., & Marshak, A. L. (1995). The interpretation of spectral vegetation indexes. IEEE Transactions on Geoscience and Remote Sensing, 33 (2), 481– 486. 10.1109/TGRS.1995.8746029

Nathan, R., Getz, W. M., Revilla, E., Holyoak, M., Kadmon, R., Saltz, D., & Smouse, P. E. (2008). A movement ecology paradigm for unifying organismal movement research. Proceedings of the National Academy of Sciences of the United States of America, 105 (49), 19052–19059. 10.1073/pnas.0800375105

Nisi, A. C., Suraci, J. P., Ranc, N., Frank, L. G., Oriol-Cotterill, A., Ekwanga, S., Williams, T. M., & Wilmers, C. C. (2022). Temporal scale of habitat selection for large carnivores: Balancing energetics, risk and finding prey. The Journal of Animal Ecology, 91 (1), 182– 195. 10.1111/1365-2656.13613

Northrup, J. M., Vander Wal, E., Bonar, M., Fieberg, J., Laforge, M. P., Leclerc, M., Prokopenko, C. M., & Gerber, B. D. (2022). Conceptual and methodological advances in habitat-selection modeling: Guidelines for ecology and evolution. Ecological Applications, 32 (1), e02470. 10.1002/eap.2470

Oliveira-Santos, L. G. R., Forester, J. D., Piovezan, U., Tomas, W. M., & Fernandez, F. A. S. (2016). Incorporating animal spatial memory in step selection functions. The Journal of Animal Ecology, 85 (2), 516–524. 10.1111/1365-2656.12485

Osipova, L., Okello, M. M., Njumbi, S. J., Ngene, S., Western, D., Hayward, M. W., & Balkenhol, N. (2019). Using step-selection functions to model landscape connectivity for african elephants: Accounting for variability across individuals and seasons. Animal Conservation, 22 (1), 35–48. 10.1111/acv.12432

Ouled-Cheikh, J., Sanpera, C., Bécares, J., Arcos, J. M., Carrasco, J. L., & Ramírez, F. (2020). Spatiotemporal analyses of tracking data reveal fine-scale, daily cycles in seabird–fisheries interactions. ICES Journal of Marine Science, 77 (7-8), 2508–2517. 10.1093/icesjms/fsaa098

Palmer, M. S., Gaynor, K. M., Becker, J. A., Abraham, J. O., Mumma, M. A., & Pringle, R. M. (2022). Dynamic landscapes of fear: Understanding spatiotemporal risk. Trends in Ecology & Evolution, 37 (10), 911–925. 10.1016/j.tree.2022.06.007

Parsons, M. B., Gillespie, T. R., Lonsdorf, E. V., Travis, D., Lipende, I., Gilagiza, B., Kamenya, S., Pintea, L., & Vazquez-Prokopec, G. M. (2014). Global positioning system data-loggers: A tool to quantify fine-scale movement of domestic animals to evaluate potential for zoonotic transmission to an endangered wildlife population. PloS one, 9 (11), e110984. 10.1371/journal.pone.0110984

Patterson, C. R., Lustig, A., Seddon, P. J., Wilson, D. J., & van Heezik, Y. (2024). Eradicating an invasive mammal requires local elimination and reduced reinvasion from an urban source population. Ecological Applications, e2949. 10.1002/eap.2949

Petty, A. M., Werner, P. A., Lehmann, C. E. R., Riley, J. E., Banfai, D. S., & Elliott, L. P. (2007). Savanna responses to feral buffalo in kakadu national park, australia. Ecological Monographs, 77 (3), 441–463. 10.1890/06-1599.1

Picardi, S., Coates, P., Kolar, J., O’Neil, S., Mathews, S., & Dahlgren, D. (2022). Behavioural state-dependent habitat selection and implications for animal translocations. The Journal of Applied Ecology, 59 (2), 624–635. 10.1111/1365-2664.14080

Picardi, S., Ranc, N., Smith, B. J., Coates, P. S., Mathews, S. R., & Dahlgren, D. K. (2021). Individual variation in temporal dynamics of post-release habitat selection. Frontiers in Conservation Science, 2. 10.3389/fcosc.2021.703906

Pohle, J., Signer, J., Eccard, J. A., Dammhahn, M., & Schlägel, U. E. (2024). How to account for behavioral states in step-selection analysis: A model comparison. PeerJ, 12, e16509. 10.7717/peerj.16509

Potts, J. R., Bastille-Rousseau, G., Murray, D. L., Schaefer, J. A., & Lewis, M. A. (2014). Predicting local and non-local effects of resources on animal space use using a mechanistic step selection model. Methods in Ecology and Evolution, 5 (3), 253–262. 10.1111/2041-210X.12150

Potts, J. R., & Börger, L. (2023). How to scale up from animal movement decisions to spatiotemporal patterns: An approach via step selection. The Journal of Animal Ecology, 92 (1), 16–29. 10.1111/1365-2656.13832

Potts, J. R., Börger, L., Strickland, B. K., & Street, G. M. (2022). Assessing the predictive power of step selection functions: How social and environmental interactions affect animal space use. Methods in Ecology and Evolution, 13 (8), 1805–1818. 10.1111/2041-210x.13904

R Core Team. (2024). R: A language and environment for statistical computing.

Rafiq, K., Jordan, N. R., Golabek, K., McNutt, J. W., Wilson, A., & Abrahms, B. (2023). Increasing ambient temperatures trigger shifts in activity patterns and temporal partitioning in a large carnivore guild. Proceedings. Biological sciences / The Royal Society, 290 (2010), 20231938. 10.1098/rspb.2023.1938

Reed, B. C., Brown, J. F., VanderZee, D., Loveland, T. R., Merchant, J. W., & Ohlen, D. O. (1994). Measuring phenological variability from satellite imagery. Journal of Vegetation Science, 5 (5), 703–714. 10.2307/3235884

Rheault, H., Anderson, C. R., Jr, Bonar, M., Marrotte, R. R., Ross, T. R., Wittemyer, G., & Northrup, J. M. (2021). Some memories never fade: Inferring Multi-Scale memory effects on habitat selection of a migratory ungulate using Step-Selection functions. Frontiers in Ecology and Evolution, 9. 10.3389/fevo.2021.702818

Richter, L., Balkenhol, N., Raab, C., Reinecke, H., Meißner, M., Herzog, S., Isselstein, J., & Signer, J. (2020). So close and yet so different: The importance of considering temporal dynamics to understand habitat selection. Basic and Applied Ecology, 43, 99–109. 10.1016/j.baae.2020.02.002

Rue, H., Martino, S., & Chopin, N. (2009). Approximate bayesian inference for latent gaussian models by using integrated nested laplace approximations. Journal of the Royal Statistical Society. Series B, Statistical methodology, 71 (2), 319–392. 10.1111/j.1467-9868.2008.00700.x

Schlägel, U. E., & Lewis, M. A. (2014). Detecting effects of spatial memory and dynamic information on animal movement decisions. Methods in Ecology and Evolution, 5 (11), 1236–1246. 10.1111/2041-210X.12284

Sells, S. N., Costello, C. M., Lukacs, P. M., Roberts, L. L., & Vinks, M. A. (2023). Predicted connectivity pathways between grizzly bear ecosystems in western montana. Biological Conservation, 284, 110199. 10.1016/j.biocon.2023.110199

Signer, J., Fieberg, J., Reineking, B., Schlägel, U., Smith, B., Balkenhol, N., & Avgar, T. (2023). Simulating animal space use from fitted integrated Step-Selection functions (iSSF). Methods in Ecology and Evolution. 10.1111/2041-210x.14263

Signer, J., Fieberg, J., & Avgar, T. (2017). Estimating utilization distributions from fitted step-selection functions. Ecosphere, 8 (4), e01771. 10.1002/ecs2.1771

Signer, J., Fieberg, J., & Avgar, T. (2019). Animal movement tools (amt): R package for managing tracking data and conducting habitat selection analyses. Ecology and Evolution, 9 (2), 880– 890. 10.1002/ece3.4823

Skeat, A. J., East, T. J., & Corbett, L. K. (1996). Impact of feral water buffalo. In C. M. Finlayson & I. Von Oertzen (Eds.), Landscape and vegetation ecology of the kakadu region, northern australia (pp. 155–177). Springer Netherlands. 10.1007/978-94-009-0133-9\_8

Sloane, D. R., Ens, E., Wunungmurra, Y., Mununggurr, L., Falk, A., Wunungmurra, R., Gumana, G., Towler, G., Preece, D., & Yirralka Rangers. (2024). Can exclusion of feral ecosystem engineers improve coastal floodplain resilience to climate change? insight from a case study in north east arnhem land, australia. Environmental Management. 10.1007/s00267-024-01940-2

Street, G. M., Fieberg, J., Rodgers, A. R., Carstensen, M., Moen, R., Moore, S. A., Windels, S. K., & Forester, J. D. (2016). Habitat functional response mitigates reduced foraging opportunity: Implications for animal fitness and space use. Landscape Ecology, 31 (9), 1939– 1953. 10.1007/s10980-016-0372-z

Tejero-Cantero, A., Boelts, J., Deistler, M., Lueckmann, J.-M., Durkan, C., Gonçalves, P., Greenberg, D., & Macke, J. (2020). Sbi: A toolkit for simulation-based inference. Journal of Open Source Software, 5 (52), 2505. 10.21105/joss.02505

Thaker, M., Gupte, P. R., Prins, H. H. T., Slotow, R., & Vanak, A. T. (2019). Fine-Scale tracking of ambient temperature and movement reveals shuttling behavior of elephants to water. Frontiers in Ecology and Evolution, 7. 10.3389/fevo.2019.00004

Thurfjell, H., Ciuti, S., & Boyce, M. (2014). Applications of step-selection functions in ecology and conservation. Movement Ecology, 2 (1), 4. 10.1186/2051-3933-2-4

Toro-Cardona, F. A., Parra, J. L., & Rojas-Soto, O. R. (2023). Predicting daily activity time through ecological niche modelling and microclimatic data. The Journal of Animal Ecology. 10.1111/1365-2656.13895

Tsalyuk, M., Kilian, W., Reineking, B., & Getz, W. M. (2019). Temporal variation in resource selection of african elephants follows long-term variability in resource availability. Ecological Monographs, 89 (2), e01348. 10.1002/ecm.1348

Tulloch, D. G. (1970). Seasonal movements and distribution of the sexes in the water buffalo, bubalus bubalis, in the northern territory. Australian Journal of Zoology, 18, 399–414.

Turchin, P. (1998). *Quantitative analysis of movement: Measuring and modeling population redistribution in animals and plants*. Sinauer Associates.

Warton, D. I. (2022). Eco-Stats: Data analysis in ecology. Springer International Publishing. 10.1007/978-3-030-88443-7

Warwick-Evans, V., Atkinson, P. W., Walkington, I., & Green, J. A. (2018). Predicting the impacts of wind farms on seabirds: An individual-based model. The Journal of Applied Ecology, 55 (2), 503–515. 10.1111/1365-2664.12996

Webb, S. L., Gee, K. L., Strickland, B. K., Demarais, S., & DeYoung, R. W. (2010). Measuring Fine-Scale White-Tailed deer movements and environmental influences using GPS collars. International Journal of Ecology, 2010. 10.1155/2010/459610

Werner, P. A. (2005). Impact of feral water buffalo and fire on growth and survival of mature savanna trees: An experimental field study in kakadu national park, northern australia. Austral Ecology, 30 (6), 625–647. 10.1111/j.1442-9993.2005.01491.x

Whittington, J., Hebblewhite, M., Baron, R. W., Ford, A. T., & Paczkowski, J. (2022). Towns and trails drive carnivore movement behaviour, resource selection, and connectivity. Movement Ecology, 10 (1), 17. 10.1186/s40462-022-00318-5

Williams, B. A., Watson, J. E. M., Beyer, H. L., Grantham, H. S., Simmonds, J. S., Alvarez, S. J., Venter, O., Strassburg, B. B. N., & Runting, R. K. (2022). Global drivers of change across tropical savannah ecosystems and insights into their management and conservation. Biological Conservation, 276, 109786. 10.1016/j.biocon.2022.109786

Wood, S. N. (2011). Fast stable restricted maximum likelihood and marginal likelihood estimation of semiparametric generalized linear models. Journal of the Royal Statistical Society. Series B, Statistical methodology, 73 (1), 3–36. 10.1111/j.1467-9868.2010.00749.x

Ylitalo, A.-K., Heikkinen, J., & Kojola, I. (2021). Analysis of central place foraging behaviour of wolves using hidden markov models. Ethology, 127 (2), 145–157. 10.1111/eth.13106

## References

Bastille-Rousseau, G., & Wittemyer, G. (2019). Leveraging multidimensional heterogeneity in resource selection to define movement tactics of animals. Ecology Letters, 22 (9), 1417–1427. 10.1111/ele.13327

Bastille-Rousseau, G., & Wittemyer, G. (2022). Simple metrics to characterize inter-individual and temporal variation in habitat selection behaviour. The Journal of Animal Ecology, 91 (8), 1693–1706. 10.1111/1365-2656.13738

Dall, S. R. X., Bell, A. M., Bolnick, D. I., & Ratnieks, F. L. W. (2012). An evolutionary ecology of individual differences (A. Sih, Ed.). Ecology Letters, 15 (10), 1189–1198. 10.1111/j.1461-0248.2012.01846.x

Fieberg, J., Freeman, S., & Signer, J. (2023, September). Evaluating goodness-of-fit of animal movement models using lineups. 10.1101/2023.09.26.559591

McElreath, R. (2020). Statistical rethinking: A bayesian course with examples in R and stan. Chapman & Hall. 10.1201/9780429029608/statistical-rethinking-richard-mcelreath

Stuber, E. F., Carlson, B. S., & Jesmer, B. R. (2022). Spatial personalities: A meta-analysis of consistent individual differences in spatial behavior. Behavioral Ecology, 33 (3), 477–486. 10.1093/beheco/arab147

